# Dorsomedial prefrontal cortex links abstract planning to motor execution

**DOI:** 10.1101/2025.08.31.673392

**Authors:** Ricardo J. Alejandro, Iris Ikink, Emmanouela Foinikianaki, Clay B. Holroyd

## Abstract

Mental planning is essential for producing action sequences. Despite the contribution of planning to many everyday activities, the neurocognitive processes that map a selected plan into a concrete course of action are unclear. We asked whether planning and execution are linked by abstract task relationships that are divorced from the specifics of action implementation. Human participants underwent functional MRI while performing a task that required encoding and maintaining abstract representations in order to plan and later apply those representations to execute action sequences. Our analyses revealed that dorsomedial prefrontal cortex encoded representations during the planning period that shifted progressively from information based on perceptual input to information associated with the abstract task context. These abstract representations were then used to translate between the action plan and its execution irrespective of the perceptual and motor details, indicating that DMPFC builds and sustains an abstract representational bridge that links planning to action.

## INTRODUCTION

Mental planning is a powerful neurocognitive mechanism that enables flexible production of action sequences. For instance, imagine a driver passing a traffic sign that indicates that a roundabout is close ahead, requiring them to examine the symbols on the sign quickly in order to determine and plan the appropriate exit (Figure 1A, left). This complex planning process entails retrieving task-relevant information from memory, associating these concepts within the current task context, simulating possible behaviors, and predicting and evaluating their outcomes in terms of the task goals and subgoals, all of which often happen without the benefit of external guidance or cues (for example, because the traffic sign has already been passed). Moreover, once a plan is selected (such as taking the fifth exit; Figure 1A, left), this information needs to be maintained until the actual roundabout is reached and the planned sequence of actions executed. The implementation of the plan requires mapping its abstract goal into a concrete course of action, all the while monitoring for performance errors and unforeseen constraints (such as encountering a roundabout with an unusual shape; Figure 1A, right). Yet although planning is essential for many everyday behaviors (Rowe et al., 2001), the cognitive and neural operations that link and translate planning to execution are largely unknown.

**Figure 1.**
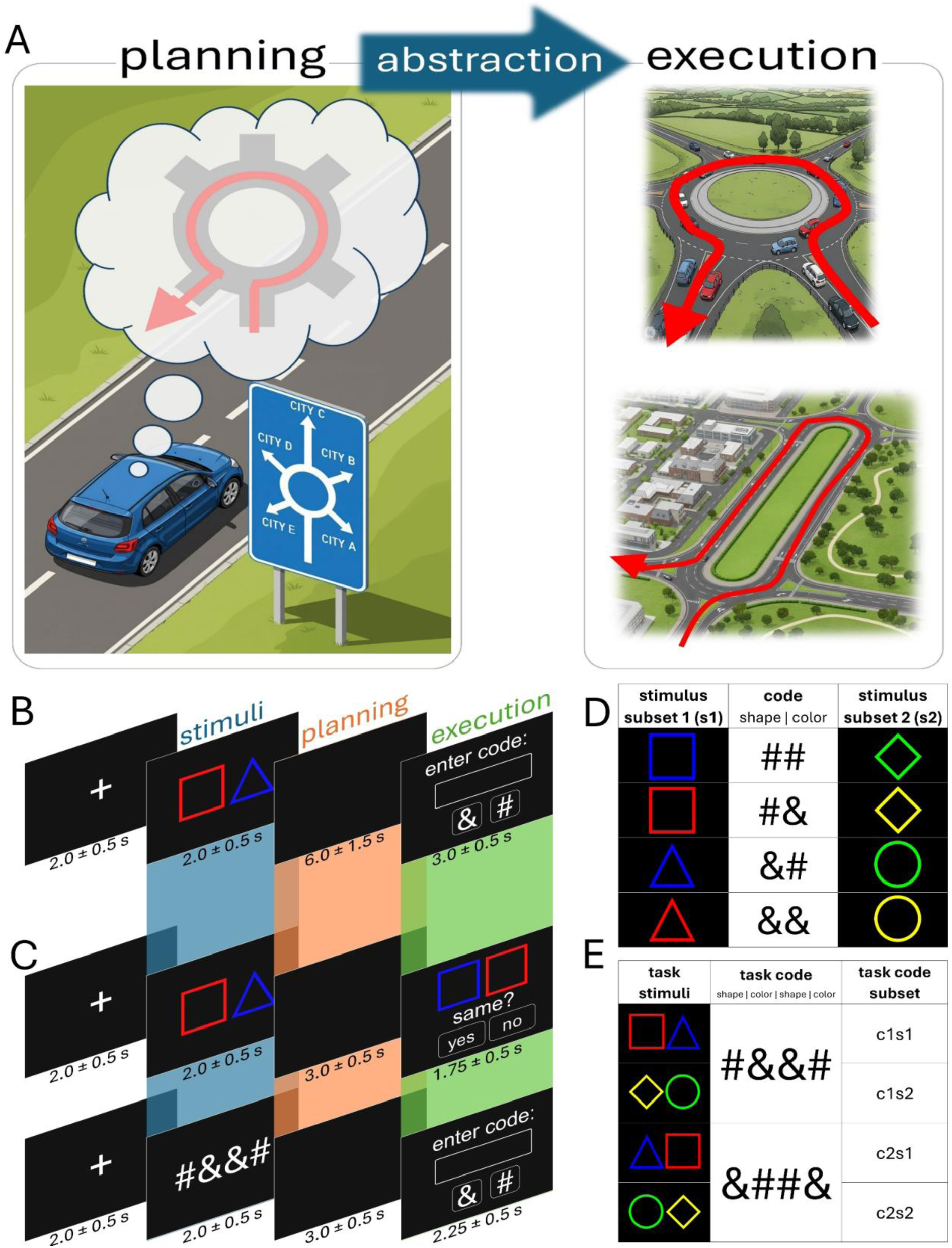
Study conceptualization and experimental paradigm(s). **A.** Illustration of the abstract link between planning and execution. The goal of a driver is to go to City E; therefore when encountering the relevant traffic sign, they determine taking the fifth exit as their plan of action. This information about prospective action must be maintained until the plan is executed because the sign has been passed and will no longer be available as a cue. Once the roundabout is reached, its physical layout might be as expected (execution, top) or unusual (execution, bottom); in both cases the driver must map their plan of action from memory to the specifics of the current environment.

Several neuroimaging studies have examined neural activity occurring during a delay or planning period between an instruction cue and its designated response (Cohen et al., 1997; Curtis & D’Esposito, 2003; Fuster & Alexander, 1971; Kubota & Niki, 1971). However, these experimental paradigms have mostly related the planning phase to a prospective motor action such as a button press, hand movement, or eye saccade. Consequently, many of these studies reported a mixture of signals related to cognitive control, action and motor preparation that are related but not specific to planning, across a distributed network of brain regions including dorsolateral-, ventrolateral-, and dorsomedial prefrontal cortex (DLPFC, VLPFC, DMPFC, respectively), anterior cingulate cortex (ACC), posterior parietal cortex (PPC), basal ganglia, cerebellum, insular cortex, hippocampus (HPC) and motor and pre-motor regions (Elsinger et al., 2006; Gallivan et al., 2011; Henderson et al., 2022; Hoshi & Tanji, 2004; Ito, 2008; Koechlin et al., 2000; Lazeron et al., 2000; Momennejad & Haynes, 2012, 2013; Rousseau et al., 2021; Ruge et al., 2010; Tanaka et al., 2021; Tosoni et al., 2014; Van Den Heuvel et al., 2003; Wagner et al., 2006; Yewbrey et al., 2023). Conversely, the execution period in these studies often reveals a combination of brain areas involved in motor and cognitive control, including DLPFC, DMPFC, PPC, cerebellum, HPC, and basal ganglia (BG) (Brandi et al., 2014; Crescentini et al., 2012; Dolfen et al., 2024; Hanakawa et al., 2008; Papitto et al., 2024; Turella et al., 2020; Unterrainer & Owen, 2006; Yewbrey & Kornysheva, 2024).

The wide variety of brain regions highlights the fact that multiple cognitive processes related to perception, cognitive control, memory, and motor control work in parallel both when planning and implementing planned behaviors as specific actions (Elsinger et al., 2006). This ubiquity of signals complicates identifying the specific contribution of each region to each of these processes (Rowe et al., 2001). Moreover, considering their reciprocal relationship, it has been noted that planning and execution are not truly separate processes (Cisek & Kalaska, 2010; Coles et al., 1985; Hommel et al., 2001). Rather, it has been proposed that planning evokes a latent, coarse representation of the action plan that is maintained in an active state until the actions are completed, in order to monitor the execution of the plan and intervene as needed (Konidaris et al., 2018). On this view, this representation would intermediate between developing a plan and implementing it, utilizing a language that is common to both task phases and that supersedes their differences. Such a high-level, abstract representation would provide the flexibility needed to map internal models to concrete courses of action while adapting the execution of the plan to external perturbations and behavioral errors (Figure 1A).

Extending this proposal, we suggest that this superordinate process, which encodes task-relevant information at an abstract level as a bridge between plans and actions, does so specifically by harnessing learned representations that generalize over extensive repertoires of tasks and task requirements (i.e., goals and constraints). Notably, the behaviors that compose a task can be organized into hierarchical structures that facilitate goal-directed behavior (Badre, 2025; Botvinick, 2008; Colin et al., 2025; Tomov et al., 2020; Wientjes & Holroyd, 2024), such that higher-order task goals and concepts modulate subordinate subgoals and explicit actions (Miller et al., 1960). Such hierarchical representations allow for existing knowledge and task properties to be generalized to new tasks (or new instances of known tasks) (Hall-McMaster et al., 2025; Lee et al., 2024; Skorstad et al., 1988; Theves et al., 2021). For instance, when learning to play the piano, a skilled guitarist does not relearn all of musical theory from scratch, but instead utilizes task-related abstractions such as notes and musical chords to integrate the commonalities across instruments (Collins & Frank, 2013; Reverberi et al., 2012; Xia & Collins, 2021). These higher-order representations enable mappings between action sequences across different domains of feature perception and motor execution (Figure 1A). Consistent with this possibility, neuroimaging evidence has revealed that hierarchically-organized plans are represented in the brain at different levels of abstraction in orbitofrontal cortex (OFC), ACC, DLPFC, DMPFC, and HPC (Balaguer et al., 2016; Brunec & Momennejad, 2022; Grossman et al., 2025; Liang et al., 2022). However, whether these areas link plans to action execution utilizing such a hierarchical encoding scheme is not clear.

Here, we identify brain areas that encode hierarchical representations for abstract planning and action execution. While undergoing functional magnetic resonance imaging (fMRI), participants were required to encode and maintain abstract representations of an upcoming action sequence and later apply those representations to execute said sequences. Crucially, this abstract context was independent of the visual stimuli and the motor actions used in the task. Following a pre-registered protocol, we trained a classifier on the multivariate fMRI activity during the planning period to discriminate different plans for sequence implementation, and then tested the classifier on the activity during the execution period. A searchlight application of this approach identified brain areas sensitive to the high-level task context that generalized over the low-level implementational details of the task. Additionally, to identify brain areas supporting abstract vs. perceptual aspects of planning, we assessed the representational similarity of brain activations with respect to two possible representational formats: abstract representations that relate different stimuli according to the task context, and perceptual representations that relate stimuli according to their shared visual features.

These analyses converged in identifying a cluster of activity overlapping DMPFC and dorsal ACC (hereafter, simply DMPFC), which suggests that this brain region maintains abstract, higher-order task representations during planning that are then used to implement the plans during their execution. In contrast, low-level visual representations related to stimulus identity were identified in visual cortex, as expected. Lastly, an exploratory analysis revealed that as the planning interval progressed, the representation during the planning period shifted from a perceptually-based format to a more abstract format.

## RESULTS

### Experimental task

Fifty participants took part in the experiment, which consisted of a main task (Figure 1B) and two functional localizer tasks (Figure 1C), while they underwent fMRI scanning. Participants were trained online beforehand during a separate session (see Methods) on task rules that required mapping colored geometric shapes into specific codes (sequence combinations of the characters “#” and “&”). On each trial of the fMRI main task (Figure 1B), the participants were shown a set of two colored geometric shapes (red and blue squares and triangles, and green and yellow diamonds and circles; Figure 1D) during a stimulus phase (blue highlight in Figure 1B), followed by a planning phase (orange highlight in Figure 1B) that required the participants to internally translate the observed sequence of colored shape stimuli into the corresponding sequential code (Figure 1E). After the planning phase, participants were shown an input field, images of two buttons labelled with the characters “#” and “&” (the relative positions of which varied at random from trial to trial), and the instruction “enter code” (execution phase, green highlight in Figure 1B). They were then required to type the corresponding (planned) 4-character code sequence as instructed on that trial (Figure 1B, D).

Hence, the plan must be encoded in a flexible format that is independent of the specific details of its execution while still directing such execution. **B.** Main task paradigm. On each trial a set of visual symbols (“stimuli”, blue shading) instructed participants to map the stimuli to the corresponding codes (predefined sequences of the characters “#” and “&”) during a planning phase (“planning”, orange shading) and subsequently type the corresponding sequence into a password prompt (“execution”, green shading). **C.** Following the main task, participants completed two functional localizer tasks (shape and code localizer). Top: functional “shape” localizer. Participants were presented with colored geometric shapes (stimuli, blue shade; same combinations as in the main task), which were instructed to be remembered during a subsequent planning phase (planning, orange shade), after which another set of colored geometric shapes were presented that either were or were not the same as the initial set during the trial, together with the question “same?” (execution, green shade). Participants were required to respond by selecting “yes” to indicate that the second set was exactly the same across all stimulus dimensions (i.e., same shapes, same colors, and same positions) as the initial set, or “no” to indicate that they were not the same in one or more of the stimulus dimensions. This localizer was designed to capture perceptual processing elements related to the main task (see Methods). Bottom: functional “code” localizer. Participants were shown an abstract code (stimuli, blue shade; same combinations of “&” and “#” as in the main task) that they were required to remember during a planning phase (planning, orange shade), after which they were prompted with a screen to execute the abstract code (execution, green shade). This localizer was designed to capture abstract processing elements related to the main task (see Methods). **D.** Visual stimuli and their corresponding abstract codes. Each geometric shape was represented by a 2-character code that was mapped to its shape (first character) and color (second character) following specific constraints. Squares and diamonds corresponded to “#”, and triangles and circles corresponded to “&”. Squares and triangles could be blue (“#”) or red (“&”), and circles and diamonds could be green (“#”) or yellow (“&”). **E.** On each trial, participants were required to execute 2-stimulus sequences of 4 button presses. To increase the amount of observations per condition, the majority of codes presented during the tasks (∼82% during the main task, ∼88% during the localizer tasks) were c1: “#&&#” and c2: “&##&” (standard trials, see Methods), which included two 2 shape-pairs per code (c1 represented by c1s1: “red square + blue triangle” or c1s2: “yellow diamond + green circle”, c2 represented by c2s1: “blue triangle + red square” or c2s2: “green circle + yellow diamond”). Our analyses focused on these “standard” trials only. During the tasks, standard trials were intermixed with catch trials that were constructed from all of the remaining valid combinations of stimuli, comprising ∼18% of the main task trials and ∼12% of the localizer tasks trials (see Methods and Table S1). The images in panel A were generated with Google Gemini and subsequently edited by the authors.

### Behavioral results

#### RT signature of chunking and benefit of planning

Because each trial required executing a four-step sequence of button presses according to each shape’s two-button sequence, we predicted that participants would encode this information hierarchically by consolidating the four-step code into two independent two-step subsequence codes (or “chunks”). To test this prediction, reaction times (RTs) to the four individual actions were compared both within-subsequence, where we predicted that the first action within each subsequence would be slower than the second (RT1 > RT2, and RT3 > RT4), and between-subsequences, where we predicted that the transition between subsequences would result in the third action to be slower than the second action (RT3 > RT2) (Eckstein & Collins, 2021; Rosenbaum et al., 1983). Additionally, behavioral evidence of planning can be seen in shorter RTs after longer planning periods (Afshar et al., 2011; Rosenbaum, 1980). We therefore predicted that the first step of the entire sequence (RT1) would be faster with increasing time for planning (namely, for planning periods of 4.5 s, 6 s, and 7.5 s; see Figure 1B and Methods).

We fitted RT data with a linear mixed-effects model (LMM) that included sequence step (1, 2, 3, 4), and planning duration (short, medium, long, given by the jittered duration of the planning phase; see Methods) as fixed effects, as well as their interactions, and random intercepts per participant. The linear mixed-effects model revealed significant main effects of sequence step (F(3, 32907) = 1418.81, p < 0.0001) and planning duration (F(2, 32908) = 1.47, p < 0.0001), and a significant interaction between sequence step and planning duration (F(6, 32907) = 6.51, p < 0.0001). Subsequent post-hoc pairwise comparisons (using Tukey’s HSD) estimated the direction of the main effects and their interaction. As shown in Figure 2A, RT1 was significantly higher than RT2 (p < 0.0001), and RT3 was significantly higher than RT4 (p < 0.0001). However, the difference between RT2 and RT3 was not statistically significant (p = 0.258). We supposed that the lack of the expected effect of hierarchical chunking (i.e., a slow-down when transitioning subsequences) could have resulted from the participants’ repeated exposure to the task both during their online behavioral training (see Methods) and during the fMRI session itself, thereby attenuating the effect. In fact, an analysis of the RT data from the online behavioral training revealed the predicted signature of hierarchical chunking (RT2 < RT3; p < 0.0001) (Figure S1), in addition to replicating the significant effects observed in the main fMRI task (RT1 > RT2, p < 0.0001); and RT3 > RT4, p < 0.0001).

**Figure 2.**
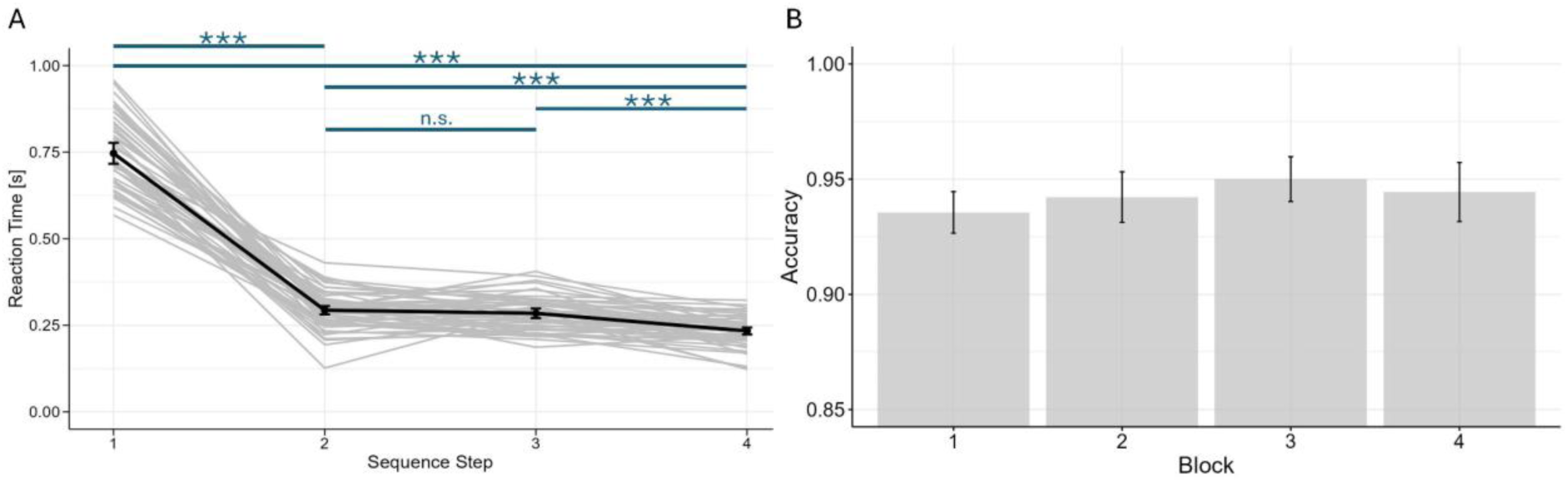
Behavioral Results. **A.** Reaction time (RT) per sequence step and post-hoc pairwise comparisons following the main effect of sequence step on RT. Bold black line represents the average RT across participants, gray lines represent individual participants. Asterisks denote statistical significance at p < 0.0001; n.s.: non-significant. **B.** Task accuracy per block; there were no significant differences across blocks (χ^2^(3) = 3.99, p = 0.262). Error bars denote the standard error of the mean (SEM) across participants.

Further, the main effect of plan duration was driven by significant differences between long and medium (7.5 vs 6 s., p = 0.0008), long and short (7.5 vs 4.5 s., p < 0.0001), and medium and short (5 vs 4.5 s., p < 0.0001) planning durations, indicating that the speed of sequence execution was influenced by the length of preparation time. Regarding the interaction between sequence step and planning duration, the main effect of planning duration on RT was only significant for RT1 when contrasted against all planning durations (long vs medium, p < 0.0001; long vs short, p < 0.0001; medium vs short, p < 0.0001). All pairwise differences for RT2, RT3, and RT4 were not statistically significant (Figure S2). This result confirmed that participants used the planning period to prepare their responses, beginning sequence execution more quickly when provided with longer periods to plan their response.

Average accuracy across participants (Figure 2B) was high (M = 0.94; SD = 0.08), with no significant differences across blocks (as revealed by a Kruskal-Wallis test: χ^2^(3) = 3.99, *p* = 0.262), as was expected due to their extensive training (see Methods).

### Neuroimaging results

#### DMPFC, DLPFC, and basal ganglia encode abstract information during planning that is used during execution

We sought to identify brain areas that mediated between superordinate, abstract sequential codes during planning and execution. To this end, following our pre-registered analysis plan we applied Multi-Voxel Pattern Analysis (MVPA) (Norman et al., 2006) to conduct a searchlight cross-decoding analysis on the multivariate fMRI data (see Figure S3 for results in pre-determined ROIs). The searchlight technique systematically inspects a spherical subset (i.e., a “searchlight”) of data from the entire brain volume that is iteratively centered on every voxel in the volume. The voxel values within each searchlight are used by a classifier to discriminate between the conditions of interest, assigning the classification accuracy score to the voxel at the center of the searchlight. This process is repeated for every voxel location within the volume, resulting in a decoding accuracy map of the entire brain (Kriegeskorte et al., 2006). During decoding, part of the data serves to train the classifier, and the rest is used for testing, which gives an estimate of the predictability of new data points based on the training set. Here, we operationalized “abstraction” as the generalization performance of a multivariate decoder on conditions not used for training (Bernardi et al., 2020). We trained the classifier to discriminate between abstract representations during the planning phase and tested it during the execution phase to predict the activity of the same representations. We hypothesized that if this abstract context connects both task phases (planning and execution), then the (abstract) information shared across phases would enable prediction of one phase from the other, regardless of the low-level (non-abstract) components of the action plan (i.e., the visual stimuli and motor responses).

Specifically, we trained a support vector machine (SVM) on the brain activity patterns (i.e., the beta value estimates, see Methods) during the planning phase of one of the stimulus subsets of each code (e.g., planning c1s1 and c2s1; Figure 3A), and used it to classify activity patterns during execution of the left-out stimuli of each code (e.g., executing c1s2 and c2s2; Figure 3A). Importantly, the task design and analysis incorporated multiple features that controlled for potential stimulus, motor, and collinearity confounds. First, the design controlled for stimulus confounds by 1) showing only a blank screen during the planning period, and 2) training the classifier on activity patterns of one set of visual stimuli (e.g., red square and blue triangle) and testing the classifier on the activity patterns of different visual stimuli that mapped to the same code (e.g., yellow diamond and green circle; Figure 3A).

**Figure 3.**
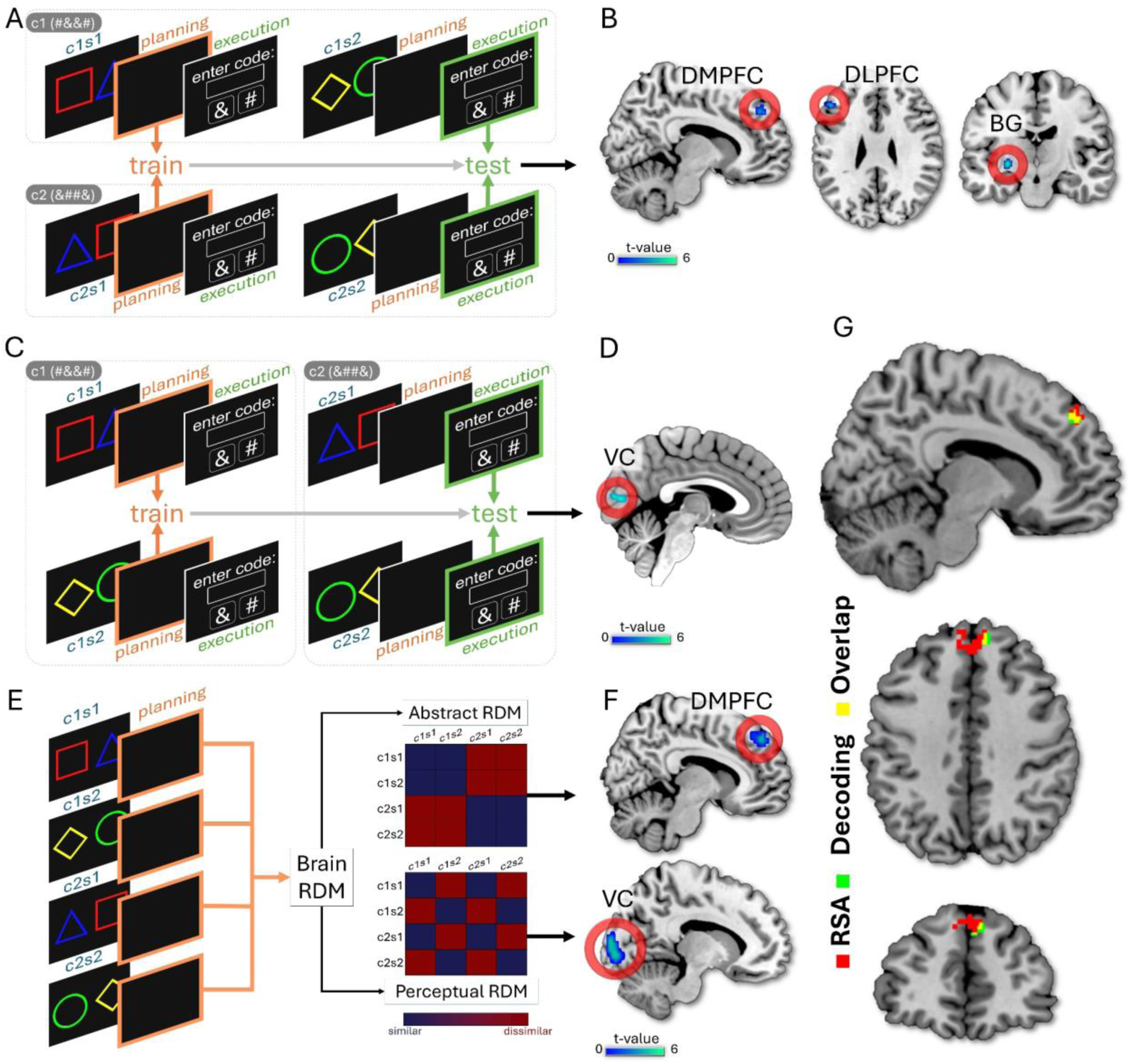
MVPA procedures and results. **A.** Pre-registered searchlight cross-decoding analysis. We trained a SVM on the brain activity patterns during the planning phase of one of the stimulus subsets of each code (e.g., planning of c1s1 and c2s1, orange highlights), and tested this classifier on brain activity patterns during execution of the left-out stimulus subsets of each code (e.g., execution of c1s2 and c2s2, green highlights; note that the position of the buttons for “&” and “#” were pseudo-randomized across trials). Training and testing took place on data from different visual stimuli (which mapped to the same abstract context), on different trial phases (planning versus execution), on different trials (stimuli for training appeared on different trials than stimuli for testing), and on different functional runs (via leave-one-run-out cross-validation). **B.** Searchlight cross-decoding results. Three clusters were identified where execution activity was decoded from planning activity at above-chance level (p_unc_ < 0.001): DMPFC (including DACC; peak at x = 10; y = 46; z = 40), DLPFC (peak at x = −48; y = 30; z = 30), and BG (putamen and pallidum, peak at x = −26; y = −14; z = −4). **C.** Control searchlight cross-decoding analysis. An SVM was trained on the brain activity patterns during the planning phase of the visual sets of each code (e.g., planning of c1s1 and c1s2, orange highlights), and tested on brain activity patterns during execution of the visual sets of the other code (e.g., execution of c2s1 and c2s2, green highlights). Training and testing took place on data from different codes (which shared high visual similarities), on different trial phases (planning versus execution), on different trials (stimuli for training appeared on different trials than stimuli for testing), and on different functional runs (via leave-one-run-out cross-validation). **D.** Control searchlight cross-decoding results. One cluster was identified where execution was decoded from planning at above-chance level (p_unc_ < 0.001): VC (peak at x = 4; y = −76; z = 10). **E.** Pre-registered searchlight representational similarity analysis (RSA). On brain activations only from the planning phase, we contrasted each participant’s “brain-RDM” (all task conditions correlated against each other: c1s1 VS c1s2 VS c2s1 VS c2s2) against two candidate “hypothesis-RDMs”: an abstract RDM was designed to capture abstract representations (representational distances depended on whether stimuli belong to within- or between code contexts), and a perceptual RDM which was designed to depict low-level visually-driven representations (distances depended on how visually similar the stimuli are). **F.** Searchlight RSA results. Top: One cluster was identified with significant similarity (p_FWE_ = 0.014) between the brain RDM and the abstract RDM, located in DMPFC (including DACC, peak at x = 3; y = 38; z = 44). Bottom: One cluster was identified with significant similarity (p_FWE_ < 0.001) between the brain RDM and the perceptual RDM, located in VC (peak at x = −18; y = −91; z = 1). **G.** Overlay of the significant clusters found by the two multivariate approaches. The same DMPFC region was found significantly active by cross-decoding planning versus execution (neural data from different trial phases) and by comparing the representational similarity during planning (neural data only from the planning phase) against hypothesis-RDMs. SVM: support vector machine; DACC: dorsal anterior cingulate cortex; DMPFC: dorsomedial prefrontal cortex; DLPFC: dorsolateral prefrontal cortex; BG: basal ganglia; VC: visual cortex.

Second, the design controlled for motor-related confounds by providing participants with the button-to-screen mappings (i.e., whether the left or the right button corresponded to the “#” and “&” symbols) only during the execution phase of the trial, which varied randomly from trial-to-trial. Third, the design incorporated several features that controlled for potential collinearity confounds between the training and testing data. In particular, because the training and testing occurred both on different trials and on different conditions (see above, and Methods), the hemodynamic response was temporally dissociated across adjacent events (Huettel, 2012; Kruggel & Von Cramon, 1999); as a consequence, the hemodynamic response to planning could not contaminate the neural representations during execution. Finally, the jittered duration of the planning phase (4.5, 6, or 7.5 s, see Methods) additionally ensured that the temporally-adjacent planning and execution regressors were sufficiently decorrelated.

All together, this MVPA procedure trained and tested the data for different conditions (different visual stimuli belonging to the same abstract context), different trial phases (associated with planning versus execution), different trials (because the stimulus for training occurred on a different trial than the stimulus for testing), and different functional runs (via leave-one-run-out cross-validation). To confirm that these measures were effective at minimizing collinearity between our measures of the planning and execution phase, we assessed two parameters that test for collinearity: pairwise regressor correlations (Josephs et al., 1997; Mumford et al., 2015) and the variance inflation factor (Mumford et al., 2025; O’brien, 2007). We found that the pairwise regressor correlations were low across the training and testing conditions (mean = −0.07, SD = 0.017; Figure S3, left), indicating that the hemodynamic responses associated with each condition were statistically separable. We further found that the variance inflation factor (VIF (Mumford et al., 2015, 2025), see Methods) was minimal for all regressors of interest across all participants (mean = 1.14, SD = 0.03; Figure S3, right), indicating that the regressors of interest were free of dependences from each other. In sum, these experimental and statistical controls ensured that our findings reflect high-level task representations rather than sensorimotor overlap or temporal dependency of the hemodynamic response.

This searchlight cross-decoding analysis yielded no significant clusters after family-wise error (FWE) rate multiple comparison correction (p_FWE_ = 0.05). However, an exploratory analysis using a less restrictive, uncorrected threshold of (p_unc_ < 0.001) identified three clusters of activity in which the representations formed during planning could be decoded at an above chance level based on the representations applied during execution (Figure 3B). One cluster encompassed DMPFC (including part of DACC, Figure 3B, left; peak at x = 10; y = 46; z = 40), another cluster was located in DLPFC (Figure 3B, middle; peak at x = −48; y = 30; z = 30) and the third cluster centered on a region of the BG (including the putamen and pallidum, Figure 3B, right; peak at x = −26; y = −14; z = −4).

#### Abstract decoding differs from perceptual decoding

To verify that these findings did not simply result from the visual similarities across different stimuli or codes, we conducted a control analysis (not pre-registered) by training an SVM classifier on the planning-related activations of the stimuli that mapped to the same abstract code (e.g., planning c1s1 and c1s2; Figure 3C) and tested it on the execution-related brain activity of the stimuli that mapped to the other code (e.g., executing c2s1 and c2s2; Figure 3C). Because each pair of stimuli denoted both codes, and thus shared a high degree of visual similarity, we hypothesized that application of the searchlight SVM cross-decoder would predominantly reveal activity clusters near visual areas. As expected, this analysis identified one significant cluster located in visual cortex (VC; Figure 3D, peak at x = 4; y = −76; z = 10).

#### DMPFC represents abstract information during planning

Via the previous multivariate decoding analysis we tested whether abstract representations of plans formed during plan preparation could be subsequently decoded from the same brain areas during the execution of those plans. Next, we investigated the abstract representational profile expressed during planning that links plans to actions, independently of the actual execution of those plans. An abstracted plan could be expressed either as a representation that generalizes across different stimuli (such that different explicit action sequences are represented by the same pattern of activations), or as individual representations that map to each action sequence. Additionally, if the planning information is based on the perceptual features (instead of on the abstract links) of the task elements, then the task concepts should relate to each other by their visual similarities. Here we focused on the abstraction that subserves generalization. We hypothesized that if the planning information is indeed represented in an abstract format, then the task concepts should generalize across their higher-level relationships.

To test these predictions, we performed a pre-registered searchlight representational similarity analysis (RSA) on brain activations recorded during the planning phase (Figure 3E). We constructed two hypothesis-driven RDMs that contrasted all the task conditions (standard trials) against each other (c1s1 VS c1s2 VS c2s1 VS c2s2): an Abstract RDM that tested for high-level, abstract relationships between plans (where the distances depended on whether the stimuli mapped to the same or different code contexts), and a Perceptual RDM that tested for low-level, visually-driven relationships between plans (where the distances depended on the visual similarity of the stimuli) (Figure 3E). We expected this hypothesis-driven RSA-searchlight to identify clusters that overlapped with the areas identified from the decoding-searchlight analysis.

The searchlight-RSA analysis yielded a single significant (p_FWE_ = 0.014) cluster representing abstract planning information in DMPFC (including parts of DACC, Figure 3F, top; peak at x = 3; y = 38; z = 44). This cluster highly overlapped the DMPFC cluster identified by the decoding analysis (Figure 3G). Likewise, the analysis yielded a single significant (p_FWE_ < 0.001) cluster representing perceptual information in VC (Figure 3F, bottom; peak at x = −18; y = −91; z = 1), which also overlapped with the VC cluster identified by the decoding analysis.

#### Shifts of representational formats during planning

Next, we investigated the temporal evolution of the abstract representations during the planning period. Toward this end, we employed two functional localizer tasks that were performed after the main fMRI task during the same functional imaging session (see Methods). The “shape” localizer (Figure 1C, top; Figure 4A, top) captured variance in the main task explained by visual processing and visual memory (such as mentally recalling the visual shapes during the planning phase). By contrast, the “code” localizer (Figure 1C, bottom; Figure 4A, bottom) captured variance in the main task explained by abstract processing alone (i.e., the abstract codes independent of the properties of their stimulus representations). We expected stronger perceptual contributions to neural activity at the beginning of the planning phase (due to visual reactivation of stimuli), resulting in greater correspondence with the shape localizer. By contrast, we expected stronger abstract contributions to neural activity at the end of the planning phase (following the translation of the instructive cues into their respective abstract codes), resulting in greater correspondence with the code localizer (Figure 4B).

**Figure 4.**
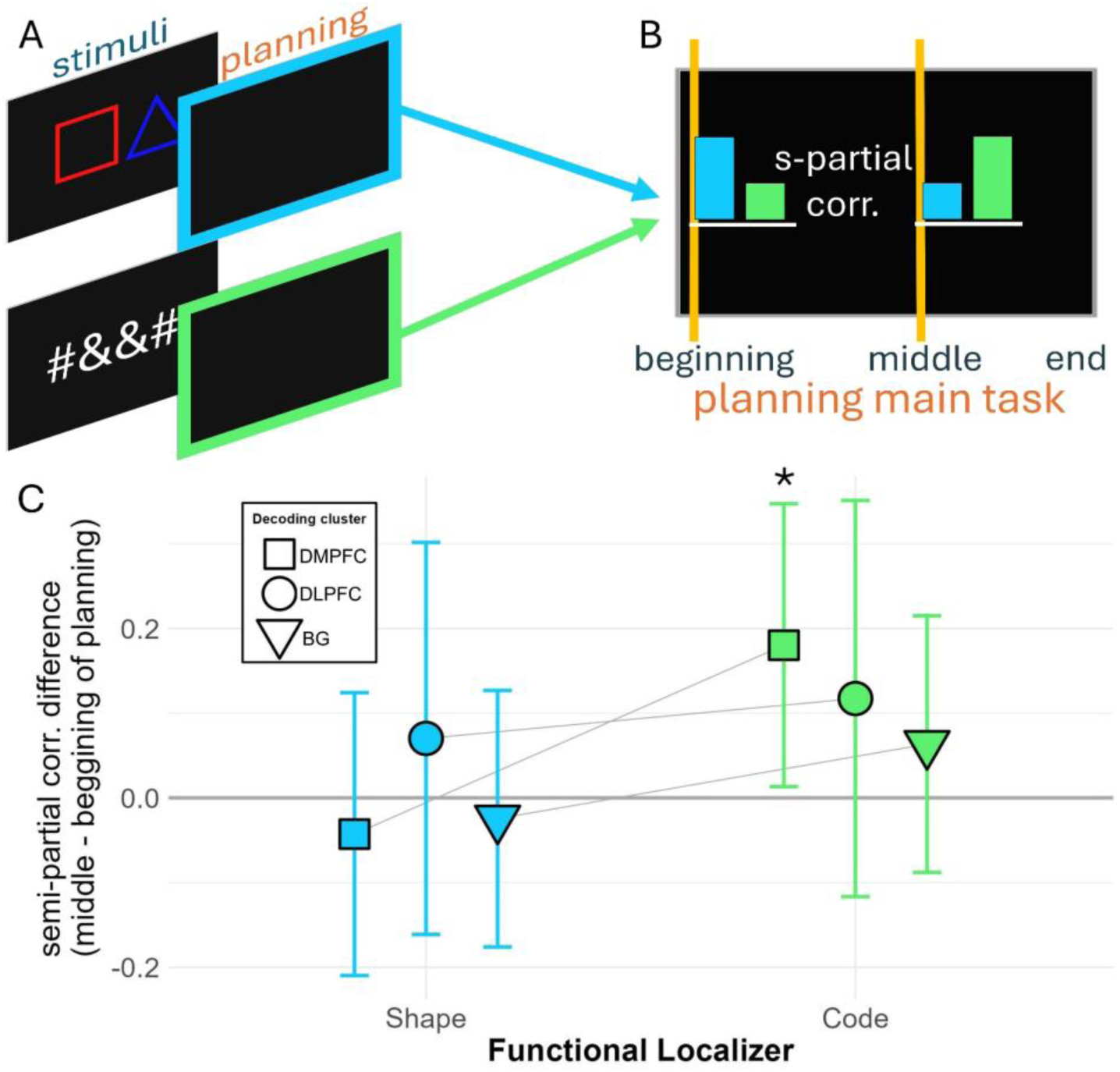
Functional localizer tasks and results. **A.** Illustration of the semi-partial correlation analysis between the planning phase of the main fMRI task and the planning phase of each of the functional localizer tasks. We correlated the patterns of activity at the beginning and middle time points during the main task with the activity patterns during the planning phase of the shape localizer (blue frame, intended to capture the variance driven by visual processing and visual memory alone) and the code localizer (green frame, intended to capture the variance driven by abstract processing alone). **B.** Predicted planning representational dynamics during the main task. We hypothesized that the beginning planning phase would be more correlated with the shape localizer, while the middle planning phase would be more correlated with the code localizer (see main text). **C.** Semi-partial correlations of cluster activity with functional localizers. The x-axis indicates the representational format (shape/perceptual vs code/abstract) during the planning period. The y-axis indicates the difference between semi-partial correlations at the middle and the beginning of the planning phase, with positive or negative numbers signaling an increase or decrease in the respective representational format (shape/perceptual vs code/abstract), respectively, from the beginning to the middle of the planning phase. Activity in the DMPFC cluster (multivariate cross-decoding analysis, see Results) was significantly more correlated with the code localizer (green square) at the middle of the planning phase, indicating an enhanced abstract representational format late in the planning period, in contrast to the correlation with the shape localizer, which did not change during the planning phase. Activity in the other brain areas (DMPFC and BG) did not change significantly during planning for either of the representational formats. Asterisk denotes statistical significance at p < 0.05.

To do so, we inspected the differential contribution of the perceptual vs abstract representations at different points in time (beginning and middle) during the planning phase in the clusters obtained during the cross-decoding analysis (DMPFC, DLPFC, and BG; Figure 3B). As pre-registered, semi-partial correlations were estimated between the patterns of activity during planning in the main task, and the patterns of activity during each of the localizer tasks (Figure 4A, B), for each of these clusters. We then tested effects of time point (beginning, middle) and localizer (shape, code) on the semi-partial correlation estimates, separately for each cluster using a two-way repeated-measures linear mixed-effects model with time point and localizer as fixed effects, as well as their interaction, and a random intercept for subject. For inference, Type III F-tests using Satterthwaite degrees of freedom were employed. This analysis returned a significant interaction between time point and localizer in DMPFC (F(1,147) = 5.98, p = 0.015). Pairwise post-hoc comparisons showed that the interaction was driven by a significant difference between the middle and beginning time points (p = 0.028) for the code localizer in DMPFC but not for the shape localizer (Figure 4C). All of the other effects across all three clusters were not statistically significant (Figure 4C).

#### Multi-dimensional scaling of planning representations

To visualize and compare the representational geometry associated with planning conditions across clusters of interest (DMPFC, DLPFC, BG), we applied multidimensional scaling (MDS (Kruskal, 1964)) to embed the planning-related RDMs (see above) into a low-dimensional space (Figure 5; analysis not pre-registered).

**Figure 5.**
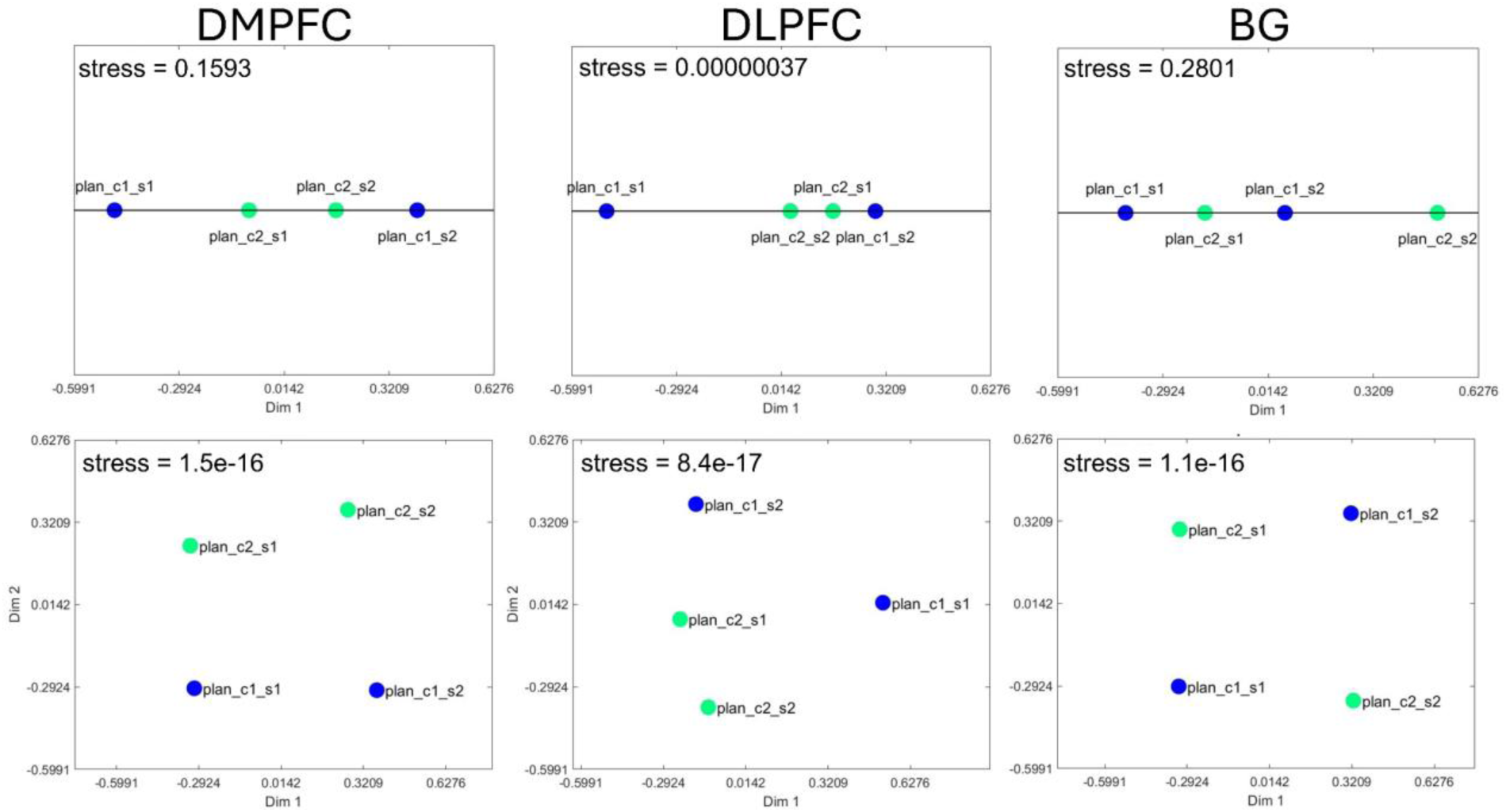
Multi-dimensional scaling (MDS) of the planning representational space. The DMPFC (left column) represents plans of upcoming actions with a two-dimensional code that distinguishes between the plans (c1 and c2) while generalizing over the stimuli (s1 and s2). By contrast, the BG (right column) is unable to separate the two plans (c1 and c2) in only two dimensions, whereas the DLPFC (middle column) is intermediate. These results suggest that all three brain areas encode plans that can be decoded during their execution, but DMPFC maintains the planning information in a relatively low-dimensional, abstract representational format.

The MDS projections reveal that DMPFC (Figure 5, left column) represents the conditions using a low-dimensional, abstract code that discriminates between the plans (c1 and c2) while generalizing over the stimuli (s1 and s2), with an excellent fit (e.g., stress = 1.5e-16) when the projection is constrained to two dimensions. By contrast, representations in the BG (Figure 5, right column) do not discriminate between the plans (c1 and c2) in a two-dimensional space despite a similarly excellent fit (stress = 1.1e-16). Lastly, representations in DLPFC (Figure 5, center column) are intermediate, fitting very well with only a single dimension (stress = 3.7e-7) despite not discriminating between conditions and requiring additional dimensions to generalize over the plans.

## DISCUSSION

Mental planning has been conceptualized as an iterative process of response selection and outcome prediction for goal-directed search, which is used to prepare or simulate upcoming sequential behaviors before their actual execution (Dehaene & Changeux, 1997). This process is understood to utilize internal representations (or models) derived from prior task experience and environmental constraints (Tolman, 1948). However, the variety of potential behaviors available for simulation often renders this process computationally intractable (Mattar & Lengyel, 2022). One strategy to make the process more efficient is to generalize over related sequences of actions in order to plan at a more abstract, hierarchically higher level, thereby simplifying the problem domain (Ho et al., 2022; Ying et al., 2025) while supporting the actual implementation of the plans. To investigate this possibility, we recorded brain activity of participants as they performed a hierarchical planning task expressly designed to test these predictions. On each trial, participants were required to map visual stimuli to abstract codes, maintain that abstract information throughout a delay period, and then implement the corresponding sequence of responses in an execution period. Behavioral analyses revealed that participants effectively used the planning period to prepare their upcoming responses (Figure S2).

Further, participants leveraged the task structure during planning by encoding the task rules hierarchically (Figure S1), though this latter effect was attenuated with repeated exposure to the task (Figure 2A).

Neuroimaging analyses revealed that the hypothesized abstract context was encoded by DMPFC (together with sections of dorsal ACC), DLPFC, and BG (including the putamen and pallidum) (Figure 3B). Specifically, an MVPA classifier was trained to discriminate between fMRI data related to different abstract codes that mapped to the same stimuli during the planning phase. That classifier was then tested on fMRI data collected during the execution phase (Figure 3A). This analysis revealed that these areas instantiated the same patterns of activity when planning and executing the behavioral sequences. These correspondences allowed for the identity of each hierarchical sequence to be predicted from the neural activity patterns occurring during its execution, based on the patterns of activity that occurred when planning an equivalent action sequence from a different, symbolic code.

Hence these representations aligned according to their higher-order contexts (as specified by the higher-order relationships between the different stimuli) rather than to the stimulus and action specifics of task execution. Critically, these results were not due to the visual similarity of stimuli corresponding to the abstract codes, as confirmed by a control analysis that trained the classifier to discriminate between different stimuli that mapped to the same abstract code (Figure 3C), which revealed activity in VC (Figure 3D). Importantly, this control analysis also argues against hemodynamic carryover from planning into execution, as it did not reproduce the main decoding result associated with planning (namely, a cluster in DMPFC). These abstract representations were also unrelated to the implementational details of plan execution, because the current experimental design prevented participants from determining the specifics of sequence execution (i.e., code-to-button mappings) until the execution phase (see Methods). Furthermore, DMPFC in particular encoded planning information in an abstract format, as revealed by searchlight-RSA based on an RDM that grouped the planning phase data according to their higher-order similarity (Figure 3F, top), whereas a searchlight-RSA based on an RDM that grouped the planning phase data according to the stimuli’s visual similarity revealed activity in VC, as expected (Figure 3F, bottom). Finally, a follow-up analysis employing functional localizers revealed that the abstract representations in DMPFC increased in strength as the delay period progressed, as would be expected of a region involved in planning the response in a higher-order space (Figure 4C).

Despite the involvement of all of these neural systems in planning, our results point to a crucial distinction between the functions of DMPFC vs. BG and DLPFC. In particular, DMPFC was the only brain region that was sensitive both to the correspondence between planning and execution and to abstract processing (Figure 3G). This property suggests that DMPFC encodes an abstract representation of a forthcoming behavioral sequence that is maintained until used, namely, during the execution of the sequence. By contrast, DLPFC and BG were identified by the searchlight cross-decoding analysis but not by the searchlight-RSA, which suggests that although these areas worked in tandem with the DMPFC to plan and produce the behavioral sequences, that information was most efficiently encoded using a low-dimensional, abstract representation in DMPFC. To gain insight into this relationship we applied multidimensional scaling (MDS (Kruskal, 1964)) to the planning representations (Figure 5; not pre-registered). Importantly, this analysis revealed that DMPFC (Figure 5, left column) clearly distinguished between the upcoming plans (c1 and c2) across stimuli using as few as two dimensions (Figure 5, top left), which suggests that this region compresses planning information into a compact, low-dimensional format. By contrast, the BG (Figure 5, right column) requires more than two dimensions to separate the planning representations. The DLPFC (Figure 5, center column) failed to discriminate between the plans despite describing the data well along only a single dimension (i.e., with about half a million times less stress compared to DMPFC and BG), which suggests a representational format that is less aligned with abstract plan identity. This distinction suggests complementary functions: whereas DLPFC and BG may be more concerned with the implementational details of the plans by relying on specialized representations that facilitate decoding (Badre, 2025), the DMPFC appears to generalize across plans using a low-dimensional, abstract contextual representation that facilitates task maintenance during planning (Figure 4C) and execution (Figure 3G).

These results are supported by previous work that has highlighted DMPFC as a center for abstract, hierarchically-organized behavior and planning (Alejandro & Holroyd, 2024; Alexander & Brown, 2015; Duverne & Koechlin, 2017; Foinikianaki et al., 2025; Holroyd & Verguts, 2021; Holroyd & Yeung, 2012; Zarr & Brown, 2016). For example, one fMRI experiment found that during a variable planning interval DMPFC was involved in sustaining and “energizing” task-relevant processes (Vallesi et al., 2009). In an fMRI study where participants planned their navigation in a virtual subway network, DMPFC activity was observed to encode the task context (i.e., current subway line), and was correlated with the cost of representing a hierarchical plan (i.e., the number of subway lines to take as opposed to the number of stations to visit) (Balaguer et al., 2016). Similarly, when navigating a virtual city, information about upcoming paths was represented across a gradient of activity in anterior prefrontal cortex (APFC, including parts of DMPFC), with more anterior regions encoding places to be visited more distantly in the future (Brunec & Momennejad, 2022). Dorsal ACC activity was also observed to encode hierarchical representations in a sequential task that emulated making different beverages (coffee vs. tea), as these representations were predicted by the activation states of a recurrent neural network (RNN) model of the task (Holroyd et al., 2018). More recently, a reanalysis of these data incorporating goal states into the RNN model demonstrated that more rostral regions of DMPFC (rostral ACC) encoded the goal-related representations in the task (Colin et al., 2025).

Further, our results align with a model-based hierarchical reinforcement learning (MB-HRL) framework that proposes that dorsal ACC is responsible for selecting and maintaining high-level task options (goals), where DLPFC follows by exerting top-down control over BG to facilitate the execution of the specific actions that implement the selected option (Holroyd & Verguts, 2021). Beyond maintaining such currently executed behaviors, we found that DMPFC prospectively maintains an abstract representation of the task context during planning and uses that representation to guide the execution of upcoming behavior.

Note that although the RSA and decoding analysis provide converging evidence on the role of DMPFC in planning, the decoding results (but not the RSA results) were determined using an uncorrected threshold and therefore should be considered exploratory. Nevertheless, the DLPFC and BG observations are consistent with a wide neuroimaging literature on this system. The prefrontal cortex (PFC) and BG are well-known to support model-based learning and planning (Fermin et al., 2016; Miller & Buschman, 2008), with the PFC constructing internal models of entire behaviors, simulating them, and directing their execution (Simon & Daw, 2011; Tricomi et al., 2009), and the BG learning action values specific to task states (Kawagoe et al., 1998; Samejima et al., 2005). Further, the activity patterns observed in DLPFC are consistent with previous arguments that this region implements rule-based response selection (Bunge et al., 2002; Cole et al., 2010). In particular, DLPFC activity has been implicated when planning information is actively monitored (Vallesi et al., 2009) and processed (Crescentini et al., 2012) (as opposed to simply maintained) during a planning delay, and when rehearsing and preparing planned responses and updating evolving task goal states (Fincham et al., 2002), for instance by selecting items and actions from working memory (Sakai & Passingham, 2003). Likewise, the BG, which are highly interconnected with the frontal cortex, have been associated with rehearsal and implementation of appropriate motor and behavioral plans (Dolfen et al., 2024; Graybiel, 2000; Groenewegen, 2003; Yewbrey & Kornysheva, 2024). The DLPFC might therefore encode a set of task rules or instruction mappings (Mian et al., 2014) that are comparable during both planning and execution and are learned via BG associations. This interpretation is consistent with the role of BG in learning stimulus-association and response selection and of DLPFC in manipulating task information compliant with task rules (Abe et al., 2007; Chatham & Badre, 2015; Cohen & Frank, 2009; Sakai & Passingham, 2006; Schumacher & D’Esposito, 2002), namely by matching shapes to codes during planning (first selecting actions internally) and from codes to button presses during execution (then executing the actions externally). In so doing, DLPFC prepares the upcoming sequence of actions (Pochon, 2001; Vallesi et al., 2009) by relating this information between planning and execution (Chatham & Badre, 2015; Sakai et al., 2002; Vaidya & Badre, 2022).

Alternate accounts highlight the HPC at the forefront of sequence planning (Yewbrey & Kornysheva, 2024) and execution (Dolfen et al., 2024). Crucially, HPC activity in those previous studies was observed when sequence planning involved knowledge not only of the behavioral sequences but also the specific temporal organization of their pre-assigned motor effectors (sequence-finger associations for execution), something that our task precluded by design. Considering the major role of HPC in the construction and maintenance of highly detailed memory episodes on the one hand, and the role of DMPFC in abstract and associative knowledge on the other, it is not surprising that DMPFC (and not HPC) was here responsible for leveraging statistical regularities between task representations in order to navigate between planning and execution. Moreover, note that some planning studies have utilized a go/no-go task design that contrasts go trials (planning followed by execution) against no-go trials (planning followed by suppression of execution) (Ariani et al., 2018; Yewbrey et al., 2023). However, such contrasts aim to isolate motor preparatory signals during planning and then relate those activations to the motor activity during execution, whereas our study investigated how a plan is translated from instruction to implementation using an abstract format that is independent of any motor and perceptual representations. The no-go requirement also introduces a variety of confounds that are unrelated to planning, such as response inhibition, which encourages participants to prepare *to not respond* when they are uncertain about whether they will be prompted to respond (Chikazoe et al., 2009). To avoid this confound, and mindful our goal to identify representations that are shared across planning and execution, we designed the task to include go-trials only, while taking specific measures to prevent collinearity in our signals of interest (see Methods and Figure S3).

These conclusions are qualified by a few caveats. First, the pre-registered behavioral chunking effect was attenuated during the main experiment (Figure 2A). However, absence of chunking does not predict absence of planning, since we observed evidence of behavioral planning when analyzing the benefit of a longer planning interval (Figure S2). Furthermore, the chunking effect was observed during the training session (Figure S1). Although the interpretation is admittedly post-hoc, we suggest that reduction of hierarchical chunking from training to testing reflects enhanced coding efficiency due to increased task exposure (given that the accuracy levels were comparable the training and main experiment sessions). Second, the pre-registered cross-decoding analysis failed to reach statistical significance after FWE correction, though it was significant at a more liberal threshold. Nevertheless, this result is supported by an independent RSA that identified this same DMPFC region in abstract planning, and by an independent functional localizer that identified planning-related representations in this DMPFC region. Evaluated collectively, these results point to an important role for DMPFC in planning. Third, the limited time resolution inherent to fMRI prevented a finer analysis of the dynamics of the neural representations. However, by relating independent functional localizers to specific planning time points, our analyses provide insight into how these representations evolved during planning, namely that DMPFC representations became more abstract as the planning interval progressed. Taken together, these results provide evidence for how plans are transformed into action. Moving forward, prospective neuroimaging studies utilizing imaging modalities with better time resolution (such as electroencephalography or magnetoencephalography) can clarify how abstract representations for planning are acquired and developed, both within and across trials.

## Conclusion

Planning and action operate along a continuum whereby planning allows for behaviors to be internally prepared and rehearsed, and action entails the rehearsed sequences of behavior to be implemented. These operations that translate planning to action require a common (abstract) representation that spans the transition from intention to execution. Crucially, real-world environments provide constraints that are sometimes revealed only during the execution of a behavior, and thus are difficult to plan for in advance. Hence, when driving on a roundabout, we might miss our turn if the roundabout has an unfamiliar layout despite having read the sign and planned our exit well in advance. Such abstract representations encode fundamental, high-level relational information that can be used to plan behavioral sequences and then adaptively translate the plans into specific details of implementation. Our results indicate that DMPFC plays a unique role in this hierarchical planning-to-action mechanism.

## METHODS

Experimental design, sample size, and statistical analyses were preregistered (https://doi.org/10.17605/OSF.IO/E58KT) and are described below. Deviations from the preregistration are noted as such.

### Participants

Fifty-three participants (38 females, mean age = 24.5, range = 18.8 – 37.5) took part in an online behavioral training session. Before the subsequent fMRI scanning session, the data of two participants were removed due to failure to complete the training (online training timed out), and the data of one other participant were removed due to MRI-incompatibility. Our sample for analysis therefore included 50 participants (35 females, mean age = 24.4, range = 18.8 – 37.5) who took part in both the online behavioral training and in the fMRI experiment. All participants were screened for MRI-compatibility and normal or corrected-to-normal visual acuity, were compensated for their participation, and provided informed consent before each experimental session (online and fMRI). The Ethics Committee of Ghent University Hospital reviewed and approved this study. This study was conducted in accord with the Declaration of Helsinki.

### Procedure

We conducted a two-part experiment. In the first part, participants underwent behavioral training online, which was intended for participants to learn the task stimuli and rules. For the second part, participants underwent fMRI scanning while performing a slightly modified version of the task (see Method Details) plus two additional functional localizer tasks inside an fMRI scanner. The two portions of the experiment were completed by the same set of participants.

### Behavioral training (online)

The online session comprised three increasingly difficult task training phases with two parts each: guided training (instructions + 10 practice trials) followed by task practice that required an accuracy criterion to be met (10 consecutive correct trials; same for all training stages) in order to proceed to the next training stage. The duration of the online training task depended on instruction reading time and task performance. However, the online training timed out after 120 minutes.

### fMRI scanning session

Within 24 hours after successfully completing the online session, participants came to the fMRI scanner for an imaging session that included a main task and two functional localizer tasks. Due to scanner malfunctions, 3 of the participants were rescheduled for a later date.

Prior to entering the scanner, participants practiced the task (a shortened version of the online training) for about 10 minutes on a laptop. Inside the scanner they performed the main task, which was similar to the task they learned in the preceding session (see Method Details) for four blocks of about 10 minutes each. After the four main task blocks, they performed two functional localizer tasks for about 10 minutes each. In total, the scanning session took around 65 minutes.

### Behavioral tasks

#### Online training task

Participants were instructed to decipher codes (combinations of characters “#” and “&”) from abstract stimuli (colored geometric shapes), and that they would use the index and middle fingers of their right hand to press the “J” and “K” keys to select items (the characters “#” and “&”) on the left and the right side of their screen, respectively. In general, the training started with an elementary task version of 1 stimulus with 1-character code (representing its shape), progressed to 1 stimulus with a 2-character code (representing shape + color), and finished with 2 stimuli with a 2-character code each (representing shape + color for each).

On the first training stage (training shape), participants were shown a single, randomly selected white shape (square, circle, diamond, or triangle) on a black background, and were instructed to select “#” if the shape was a square or a diamond, or “&” if the shape was a triangle or a circle. A trial ended when the correct response was chosen. They performed 10 practice trials irrespective of their accuracy. Then, in order to advance to the next training stage, they were required to correctly complete two 10-trial sequences of consecutive trials (thus meeting an accuracy criterion).

For the second training stage, a delay screen was introduced after stimulus presentation, during which the participants were required to plan the code that they would enter in the subsequent execution phase. As with the first training stage, they performed 10 practice trials, which they repeated until reaching the accuracy criterion.

For the next (third) training stage (training shape + color), the participants were presented with the same shapes as before (square, diamond, circle, triangle) but now displayed in specific colors (squares and triangles were either blue or red, circles and diamonds were either green or yellow; Figure 1C), selected at random. Consequently, the codes to be input had two dimensions (i.e., character shape and character color). Initially, the participants were presented with each colored shape and were asked to input the 2-character code, and afterwards a planning phase was introduced (similar to stage 1). Finally, a time limit was introduced for the execution phase. The participants proceeded to the next stage once they have reached the accuracy criterion.

In the final (fourth) training stage (training shape + color with 2 stimuli), the stimuli at each trial consisted of 2 shapes (a shape pair) presented together. Participants were required to input two 2-character codes one after the other (equivalent to one 4-character code). Similar as the proceeding training stages, participants were trained in blocks without a planning phase, with a planning phase, and with a time limit for execution. They proceeded to the next stage once they had reached accuracy criterion.

After participants completed the training stages, they performed the “main training task” by performing 4 blocks of the task with two shape pairs (describing two 2-character codes). This stage was conducted without an accuracy criterion. Accuracy across blocks for the main training task was high (M = 0.93, SD = 0.007, range = 0.92 – 0.94) and comparable to the main fMRI task (M = 0.94, SD = 0.007, range = 0.93 – 0.95).

#### fMRI tasks

Each imaging session consisted of a main task and two functional localizer tasks. For the main task, participants were placed inside the scanner and performed 4 blocks of around 10 minutes each of a task similar to the main training task. Each trial started with a fixation cross, followed by the presentation of the trial stimuli (colored geometric shapes). The durations of both the fixation cross and the stimuli were jittered by 2.0 ± 0.5 s. After stimulus presentation, a black screen was presented for 6.0 ± 1.5 s. (jittered), signaling the planning phase. During this period participants were required to map the stimuli (colored geometric shapes) presented on that trial to their corresponding codes (that were learned during the practice session). Finally, the execution phase was signaled by the presentation on screen of an input field, the instruction “enter code”, and two buttons labelled with the characters “#” and “&” (the relative positions of these buttons varied at random from trial to trial). The execution phase lasted 3.0 ± 0.5 s. (jittered), and participants were required to enter the codes corresponding to the trial stimuli by pressing two spatially equivalent buttons (i.e., right button to select right item on the screen) on an MRI-compatible button box using the right-hand index and middle fingers. A feedback screen was presented only for incorrect trials with a fixed duration of 0.5 s.

The duration of each of the phases (fixation cross, stimuli presentation, planning, and execution) were assigned pseudo-randomly on each trial, such that all the trials in the experiment lasted for 13 s, and all possible durations were presented in equal proportion (e.g., the planning phase lasted an equal number of times for 4.5, 6.0, or 7.5 s).

The fMRI task differed from the (online) main training task (see above) according to the relative proportions of the presented stimuli. To increase the number of observations per condition, we restricted the number of codes to be analyzed to only two (c1: “#&&#”, c2: “&##&”), each of which was represented by two 2 shape-pairs -- i.e., c1 represented by c1s1 (“red square + blue triangle”) and c1s2 (“yellow diamond + green circle”), and c2 represented by c2s1 (“blue triangle + red square”) and c2s2 (“green circle + yellow diamond”) (Figure 1E). Each main task run contained 44 trials, about 82% of which consisted of the trials of interest or “standard trials” (c1 or c2; Figure 1E), and the remaining 18% were “catch trials” (Table S1) that were randomly selected from the allowed feature combinations (i.e.., shapes in s1 combined only with shapes and colors within s1, and shapes in s2 combined only with shapes and colors within s2; Figure 1D). Standard and catch trials were presented intermixed during each task run.

After completing the main task, participants performed two functional localizer tasks (“shape” and “code” localizers) that lasted approximately 10 minutes each. The shape localizer task included 68 task trials, including 15 trials per standard-trial shape-pair (or 30 trials per each of the two standard-trial codes), and 8 (∼12%) catch trials. During each trial of the shape localizer task, participants first saw a fixation cross, followed by the presentation of a shape-pair (according to the same stimuli and constraints as in the main task). The durations of both the fixation cross and the stimulus presentations were jittered by 2.00 ± 0.50 s. Next, a black screen appeared for 3.00 ± 0.50 s (jittered across trials). During this planning phase participants were required to maintain the stimulus information in working memory. Then, a shape-pair with the same stimulus properties and constraints as in the main task (including both standard and catch trials) was presented for a duration of 1.75 ± 0.50 s. During this execution phase, participants were required to indicate whether the stimuli were identical to those presented at the start of the trial by pressing one of two buttons on a response box corresponding to the answers “yes” and “no”. The correspondence between the button location (left or right) and the response options (“yes” and “no”) was cued on the screen and was pseudo-randomized across trials such that each button corresponded to “yes” and “no” an equal proportion of times. The duration of each of the phases (fixation cross, stimulus presentation, planning, and execution) were assigned pseudo-randomly on each trial, such that all the trials in the shape localizer task lasted for 8.75 s, and all possible durations were presented in equal proportion (e.g., the planning phase lasted an equal number of times for 2.50, 3.00, or 3.50 s).

The code localizer task included 68 task trials (30 trials per each of the two standard-trial codes) and 8 (∼12%) catch trials. During each trial, participants were presented with a fixation cross (2.00 ± 0.50 s) and then an abstract code consisting of the same combinations of the symbols “&” and “#” as they appeared in the main task (2.00 ± 0.50 s), the duration of which was jittered across trials. Then a black screen was displayed for 3.00 ± 0.50 s jittered across trials, during which the participants were required to remember the observed code. Finally, a screen that was identical to the execution phase of the main task appeared for 2.25 ± 0.50 s, during which participants were required to execute a response (see above and Figure 1). The duration of each of the phases (fixation cross, stimulus presentation, planning, and execution) were assigned pseudo-randomly for each trial such that all the trials in the code localizer task lasted for 9.25 s, with all possible durations presented in equal proportions (e.g., the planning phase lasted an equal number of times for 2.50, 3.00, or 3.50 s).

For both the main task and the functional localizer tasks, the standard and catch trials were presented intermixed. The standard trial codes were constructed such that 1) each character appeared an equal number of times in each code, 2) the codes were composed of more than one symbol (excluding, for example, the stimulus “####”), and 3) the dimensions were orthogonal across codes (e.g., red in code 1 is blue in code 2). During the scanning session all of the participants performed the main task first, followed by the two localizer tasks; the order of the two localizer tasks was counterbalanced across participants.

Participants pressed the left and right buttons of the MRI-compatible button box to select the code characters (“#” and “&”) that were presented on the left and the right side of the screen. Importantly, the location of the characters on the screen was assigned pseudorandomly across trials and participants, such that either code character appeared at the same (left or right) screen location for no more than 2 consecutive trials.

### MRI Data Collection

MRI data were collected in a Siemens MAGNETOM PrismaFit 3T MRI scanner using a 64-channel head coil. For each participant we collected a high-resolution T1-weighted anatomical scan using an MPRAGE sequence (176 slices; voxel size = 1 mm isotropic; TR = 2250 ms; TE = 4.18 ms; FoV = 256 mm), and whole-brain, T2*-weighted multi-band echo-planar imaging (EPI) functional scans (54 slices per volume; TR = 1780 ms; TE = 27 ms; voxel size = 2.5 mm isotropic; FoV = 210 mm; interleaved slice acquisition).

### Behavioral data analysis

R (version 4.4.0) running under RStudio (version 2024.09.1+394) was used for all behavioral analyses. Per participant, we estimated mean reaction times (RTs) per sequence step, and accuracies with the Rmisc package. We performed a Kruskal-Wallis test to assess overall differences in accuracy across blocks.

One RT was estimated for each of the 4 steps in the sequence: RT for step 1 was measured from the onset of execution until the first button press, and RTs for steps 2–4 were measured as the intervals between successive button presses (e.g., RT2 was measured from the first to the second button press). RTs were analyzed fitting a linear mixed-effects model (LMM) using the lme4 package. The model included step (1, 2, 3, 4) and planning duration (short, medium, long) as fixed effects, as well as their interaction. To account for individual differences, we included random intercepts for subjects. We assessed the significance of the fixed effects using Type III ANOVA with Satterthwaite’s degrees of freedom approximation. For significant main effects or interactions, we conducted post-hoc pairwise comparisons of estimated marginal means with Tukey’s Honestly Significant Difference (HSD) correction implemented in the emmeans R package.

### MRI data preprocessing

Before preprocessing, raw (DICOM) MRI data were converted into BIDS-format (Gorgolewski et al., 2017) using BIDScoin 3.0.8. Results included in this manuscript come from preprocessing performed using fMRIPrep 23.2.3, a Nipype 1.8.6 (Gorgolewski et al., 2011) based tool. As recommended by the fMRIPrep developers, we report below the automatically generated outputs from preprocessing both the anatomical and the functional MRI data.

### Anatomical data preprocessing

A total of 1 T1-weighted (T1w) images were found within the input BIDS dataset. The T1w image was corrected for intensity non-uniformity (INU) with N4BiasFieldCorrection (Tustison et al., 2010), distributed with ANTs 2.5.0 (Avants et al., 2008), and used as T1w-reference throughout the workflow. The T1w-reference was then skull-stripped with a Nipype implementation of the antsBrainExtraction.sh workflow (from ANTs), using OASIS30ANTs as target template. Brain tissue segmentation of cerebrospinal fluid (CSF), white-matter (WM) and gray-matter (GM) was performed on the brain-extracted T1w using fast (Zhang et al., 2001) (FSL). Brain surfaces were reconstructed using recon-all (FreeSurfer 7.3.2), and the brain mask estimated previously was refined with a custom variation of the method to reconcile ANTs-derived and FreeSurfer-derived segmentations of the cortical gray-matter of Mindboggle (RRID:SCR_002438). Volume-based spatial normalization to one standard space (MNI152NLin2009cAsym) was performed through nonlinear registration with antsRegistration (ANTs 2.5.0), using brain-extracted versions of both T1w reference and the T1w template. The following template was selected for spatial normalization and accessed with TemplateFlow (23.1.0): ICBM 152 Nonlinear Asymmetrical template version 2009c (TemplateFlow ID: MNI152NLin2009cAsym).

### Functional data preprocessing

For each of the 6 BOLD runs found per subject (across all tasks and sessions), the following preprocessing was performed. First, a reference volume was generated, using a custom methodology of fMRIPrep, for use in head motion correction. Head-motion parameters with respect to the BOLD reference (transformation matrices, and six corresponding rotation and translation parameters) are estimated before any spatiotemporal filtering using mcflirt (Jenkinson et al., 2002) (FSL). The estimated fieldmap was then aligned with rigid-registration to the target EPI (echo-planar imaging) reference run. The field coefficients were mapped on to the reference EPI using the transform. The BOLD reference was then co-registered to the T1w reference using bbregister (FreeSurfer) which implements boundary-based registration (Greve & Fischl, 2009). Co-registration was configured with six degrees of freedom. Several confounding time-series were calculated based on the preprocessed BOLD: framewise displacement (FD), DVARS and three region-wise global signals. FD was computed using two formulations following Power (absolute sum of relative motions)(Power et al., 2014), and Jenkinson (relative root mean square displacement between affines)(Jenkinson et al., 2002). FD and DVARS are calculated for each functional run, both using their implementations in Nipype (following the definitions by Power et al.(Power et al., 2014)). The three global signals are extracted within the CSF, the WM, and the whole-brain masks. Additionally, a set of physiological regressors were extracted to allow for component-based noise correction (CompCor, Behzadi et al., 2007). Principal components are estimated after high-pass filtering the preprocessed BOLD time-series (using a discrete cosine filter with 128s cut-off) for the two CompCor variants: temporal (tCompCor) and anatomical (aCompCor). tCompCor components are then calculated from the top 2% variable voxels within the brain mask. For aCompCor, three probabilistic masks (CSF, WM and combined CSF+WM) are generated in anatomical space. The implementation differs from that of Behzadi et al. in that instead of eroding the masks by 2 pixels on BOLD space, a mask of pixels that likely contain a volume fraction of GM is subtracted from the aCompCor masks. This mask is obtained by dilating a GM mask extracted from the FreeSurfer’s aseg segmentation, and it ensures components are not extracted from voxels containing a minimal fraction of GM. Finally, these masks are resampled into BOLD space and binarized by thresholding at 0.99 (as in the original implementation). Components are also calculated separately within the WM and CSF masks. For each CompCor decomposition, the k components with the largest singular values are retained, such that the retained components’ time series are sufficient to explain 50 percent of variance across the nuisance mask (CSF, WM, combined, or temporal). The remaining components are dropped from consideration. The head-motion estimates calculated in the correction step were also placed within the corresponding confounds file. The confound time series derived from head motion estimates and global signals were expanded with the inclusion of temporal derivatives and quadratic terms for each (Satterthwaite et al., 2013). Frames that exceeded a threshold of 0.5 mm FD or 1.5 standardized DVARS were annotated as motion outliers. Additional nuisance timeseries are calculated by means of principal components analysis of the signal found within a thin band (crown) of voxels around the edge of the brain, as proposed by (Patriat et al., 2017). All resamplings can be performed with a single interpolation step by composing all the pertinent transformations (i.e. head-motion transform matrices, susceptibility distortion correction when available, and co-registrations to anatomical and output spaces). Gridded (volumetric) resamplings were performed using nitransforms, configured with cubic B-spline interpolation.

### fMRI data analysis

Neuroimaging analyses were conducted using SPM12 software (Statistical Parametric Mapping; Wellcome Department of Imaging Neuroscience, London, UK) running on Matlab (version R2023a) with customized scripts and in tandem with purpose-specific toolboxes: The Decoding Toolbox (version 3.99F (Hebart et al., 2015)) for decoding analysis, and the rsatoollbox (Nili et al., 2014) for representational similarity analysis (RSA).

To estimate voxel activity patterns via MVPA, we used multivariate decoding (Norman et al., 2006), and Representational Similarity Analysis (Kriegeskorte et al., 2008). We first ran a voxel-wise general linear model (GLM) on unsmoothed, unnormalized individual data. This GLM included regressors of interest for planning and execution phases, modelled as boxcars. Error and late trials were modelled as regressors that comprised the duration of the entire trial. Confound regressors (estimated during preprocessing) included six head motion parameters, framewise displacement, and global signal (within cerebrospinal fluid, white matter, and brain mask). All regressors were convolved with a canonical hemodynamic response function plus time and dispersion derivatives as implemented in SPM12. The resulting beta maps were used to perform our searchlight MVPA analyses. Additionally, region-of-interest analyses were performed for multivariate decoding-, representational similarity-, and semi-partial correlation analyses (Figure S3). As preregistered, we used a p-value of 0.05 to establish statistical significance, and to control for multiple comparisons where applicable we used a familywise error (FWE) cluster correction of p = 0.05 and a primary voxel-wise threshold of p < 0.001.

### Multi-Voxel Pattern Analysis (MVPA): searchlight cross-decoding of action execution from action planning

To answer our primary question of whether the brain maintains an abstract, higher-order context during planning and uses this representation when executing the plan, we conducted a one-vs-one multi-class cross-validated (leave-one-run-out) searchlight cross-classification of the executed action sequences, with classification accuracy minus chance values per voxel (accuracy maps) as the output. Beta maps from the planning phase of one shape-pair from each condition (e.g., c1s1 and c2s1) were used for training, and beta maps from the execution phase of the left-out conditions (e.g., c1s2 and c2s2) were used for testing (Figure 3A). This procedure was repeated by switching the conditions (i.e., training on the planning phases of c1s2 and c2s2, and testing on the execution phase of c1s1 and c2s1, respectively; Figure 3A). Crucially, because this procedure used one set of conditions for training and a different set of conditions for testing, the training and testing split was carried out on data taken from different trials (because different conditions took place on different trials), different trial phases (training on planning vs testing on execution), different planning conditions (because the classifier was trained on one set stimuli and tested on a different set of stimuli), and different functional runs (via leave-one-run-out cross-validation), per subject. The output of this analysis consisted of an accuracy map per subject (from planning to execution), indicating the degree to which the abstract codes generalized across task phases regardless of their low-level stimulus features and motor demands.

The analysis was run using a linear support vector machine (SVM) implemented within The Decoding Toolbox (Hebart et al., 2015) software package. Decoding was performed in native space by scanning a spherical searchlight (6 mm of radius) across the condition beta values for the entire brain to obtain an accuracy map (cross-decoding accuracy score assigned to each searchlight-centered voxel) per subject. We normalized each subject’s z-scored accuracy map into MNI space and applied spatial smoothing using a 3 mm FWHM gaussian kernel. Statistical significance was assessed at the group level by performing a one-sample t-test against the null hypothesis (chance performance level of 0.25) across participants followed by cluster-wise inference. Specifically, on the resulting map we applied a recommended voxel-level cluster-defining threshold of p < 0.001 (uncorrected) (Woo et al., 2014).

Then, cluster significance was determined using familywise error (FWE) correction at p < 0.05 at the cluster level, based on Gaussian Random Field Theory as implemented in SPM. Finally, for visualization purposes, the resulting statistical map was thresholded at a cluster size of 20 voxels.

Considering that the activity of interest for this cross-decoding analysis took place in sequential temporal periods, we took several measures to mitigate potential contamination of neural activity occurring during the execution phase by neural activity occurring during the planning phase, as well as other confounds inherent to fMRI research. First, to prevent activation related to motor preparation from occurring during the planning phase, a blank screen was shown during the planning period, followed by an image of the button-to-screen mappings during the execution period. Second, in order to mitigate against the lag of the hemodynamic response from contaminating the execution period with stimulus-related neural activity occurring during planning, the data for training (on the planning representations) and testing (on the execution representations) were taken from different trials and from different conditions (for example, training on the plan for the red square and blue triangle while testing on the execution of the yellow diamond and green circle; Figure 3A). In addition, the duration of the planning phase was jittered across trials (Figure 1B), as randomized timing of events has been shown to ameliorate temporal overlap due to the hemodynamic lag (Dale, 1999). Altogether, these measures were expected to minimize collinearity among the regressors of interest.

We verified the validity of our experimental design and analysis by assessing (not pre-registered) the 1) pairwise correlations (Josephs et al., 1997; Mumford et al., 2015) and 2) the variance inflation factor (VIF) (James et al., 2013; Mumford et al., 2015) between the regressors of interest (Figure S3, left and right, respectively). The value of the VIF for regressor *X*_*i*_ is estimated by *VIF*_*i*_ = 1⁄(1 − *R*_i_^2^), where *R*_i_^2^ is the coefficient of determination obtained by modelling *X*_*i*_ as a function of all other regressors of interest (*X*_*i*_ = β_1_*X*_1_ + ⋯ + β_*i*−1_*X*_*i*−1_ + β_*i*+1_*X*_*i*+1_ + ⋯ + β_*N*_*X*_*N*_ + ε). Since the magnitude of the VIF indicates the extent to which each regressor is expressed as a linear combination of the other regressors, it is effectively a direct measure of collinearity. Values of VIF closer to 1 reflect greater independence between regressors and hence no inflation of variance, while a typical threshold of VIF = 5 or 10 is considered to signal a collinearity problem.

As expected, the VIF was low for all regressors of interest (mean = 1.14, SD = 0.03; Figure S3, right). Further, although the pairwise correlations between planning and execution regressors from the same condition/trial were moderate (mean = 0.29, SD = 0.04; Figure S3, left), the pairwise correlations between planning and execution regressors from different conditions/trials were low (mean = −0.07, SD = 0.017; Figure S3, left). Note that this decoding analysis compared representations only across conditions and trials, never within a condition or trial. Therefore, this analysis confirms that the decoded activations between planning and execution result from shared components between those stages rather than from any lingering effects of the hemodynamic response.

### Representational Similarity Analysis (RSA): searchlight RSA of planned sequences and hypothesis RDMs

To investigate the format in which information is represented while planning the execution of a sequence, a cross-validated (leave-one-run-out) searchlight representational similarity analysis (RSA) of planned sequences was applied to the data of the planning phase of the experiment. For the data of each participant, a 6 mm radius sphere was slid across the brain image in native space, obtaining for each voxel a vector of neural activations per condition (c1s1, c1s2, c2s1, c2s2). Then, on each voxel we constructed a 4×4 brain-RDM by estimating the cross-validated Pearson (1 - Pearson) pair-wise correlation across all conditions (c1s1 VS c1s2 VS c2s1 VS c2s2). On each fold of the cross-validation process the dissimilarity estimates were averaged across runs of the training set, and this averaged value was contrasted against the left-out set. The dissimilarity values were averaged across cross-validation folds to obtain the RDM estimate per voxel. We contrasted this “brain-RDM” against two candidate “hypothesis-RDMs” (Figure 3E): an *abstract* RDM that encoded whether the stimuli mapped to within- or between code contexts, and a *perceptual* RDM that encoded the visual similarity of the stimuli. To contrast the brain-RDM against each of the hypothesis-RDMs, from each voxel’s RDM we extracted the elements above the matrix diagonal, producing a 6-element vector that was compared using Spearman’s rank correlation to an equivalent 6-element vector extracted from each hypothesis-RDM. At the group level, for each hypothesis-RDM we performed a one-sample t-test. On the resulting statistical map we applied the recommended voxel-level cluster-defining threshold of p < 0.001 (uncorrected) (Woo et al., 2014). Then, cluster significance was determined using familywise error (FWE) correction at p < 0.05 at the cluster level, based on Gaussian Random Field Theory as implemented in SPM. Finally, we report clusters with activity patterns significantly correlated with each hypothesis-RDM; for clarity, clusters smaller than 20 voxels were not considered.

### Semi-partial correlation: main task and functional localizers

We hypothesized that during the planning phase of the main task, participants initially recalled the geometric shapes and progressively mapped the presented shapes to their specific codes, establishing the correct code sequence for that trial. Accordingly, we designed two functional localizer tasks to capture each of these processes: a *shape* functional localizer (Figure 1C, top) that was sensitive to the visual properties of the stimuli, and a *code* functional localizer (Figure 1C, bottom) that required implementing the abstract codes as action sequences.

The beta maps for this analysis were estimated from voxel-wise GLMs performed independently on the main task brain data and on each of the functional localizers. For the main task, we included regressors for planning at the beginning and middle time points (the middle time point varied due to the jittered duration of the planning period, see above), modelled as delta functions. For the functional localizers, we included regressors for planning at the beginning of the planning phase. For all GLMs, error and late trials were modelled as regressors comprising the duration of the entire trial. Confound regressors (estimated during preprocessing) included six head motion parameters, framewise displacement, and global signal (within cerebrospinal fluid, white matter, and brain mask). All regressors were convolved with a canonical HRF plus time and dispersion derivatives as implemented in SPM12.

To inspect how perceptual and abstract representational formats evolved across the planning phase, we analyzed the brain activity patterns at the beginning and middle time points of the planning phase focused on the clusters obtained from the cross-decoding analysis (see Results and Figure 3) as regions of interest (ROIs), as well as additional pre-registered ROIs (see Figure S4). Following a template tracking procedure (Palenciano et al., 2023), we estimated semi-partial correlations between the patterns of activity (beta maps) of the main task in each condition during the planning period, and the patterns of activity of each of the functional localizers during the planning period (Figure 4A). We used semi-partial correlations to assess the individual/unique contribution of each representational format (e.g., abstract) with respect to the main task while partializing out the contribution of the other representational format (e.g., perceptual).

We predicted that the planning-related activity in the regions of interest would be more correlated with the shape localizer than with the code localizer at the beginning of the planning phase, whereas the opposite (higher correlation with the code localizer) would be true during the middle of the planning phase. To test these predictions we first estimated the semi-partial correlation between each localizer and the main task while controlling for the influence of the other localizer on the main task, that is, we correlated the activity patterns of one localizer (e.g., code) with the residuals obtained from regressing the activity patterns of the other localizer (e.g., shape) onto the activity patterns of the main task. We then repeated this procedure at each time point (beginning, middle) for each ROI (DMPFC, DLPFC, and BG clusters). Then, for each ROI, the semi-partial correlation estimates were analyzed via a two-way repeated measures linear mixed-effects model with fixed effects of time point (beginning, middle), localizer (shape, code), and their interaction, and subject included as a random effect (random intercept). Fixed effects were evaluated with Type III F-tests using Satterthwaite degrees of freedom. Post-hoc comparisons were tested using estimated marginal means (emmeans), with pairwise contrasts corrected for multiple comparisons using the Tukey method.

## ACKNOWLEDGMENTS

This project was supported by funding from the European Research Council (ERC) under the EU’s Horizon 2020 Research and Innovation Programme (grant agreement no. 787307).

## DATA AND CODE AVAILABILITY

Data and code will be made available upon acceptance for publication.

## SUPPLEMENTARY INFORMATION

**Table S1.**
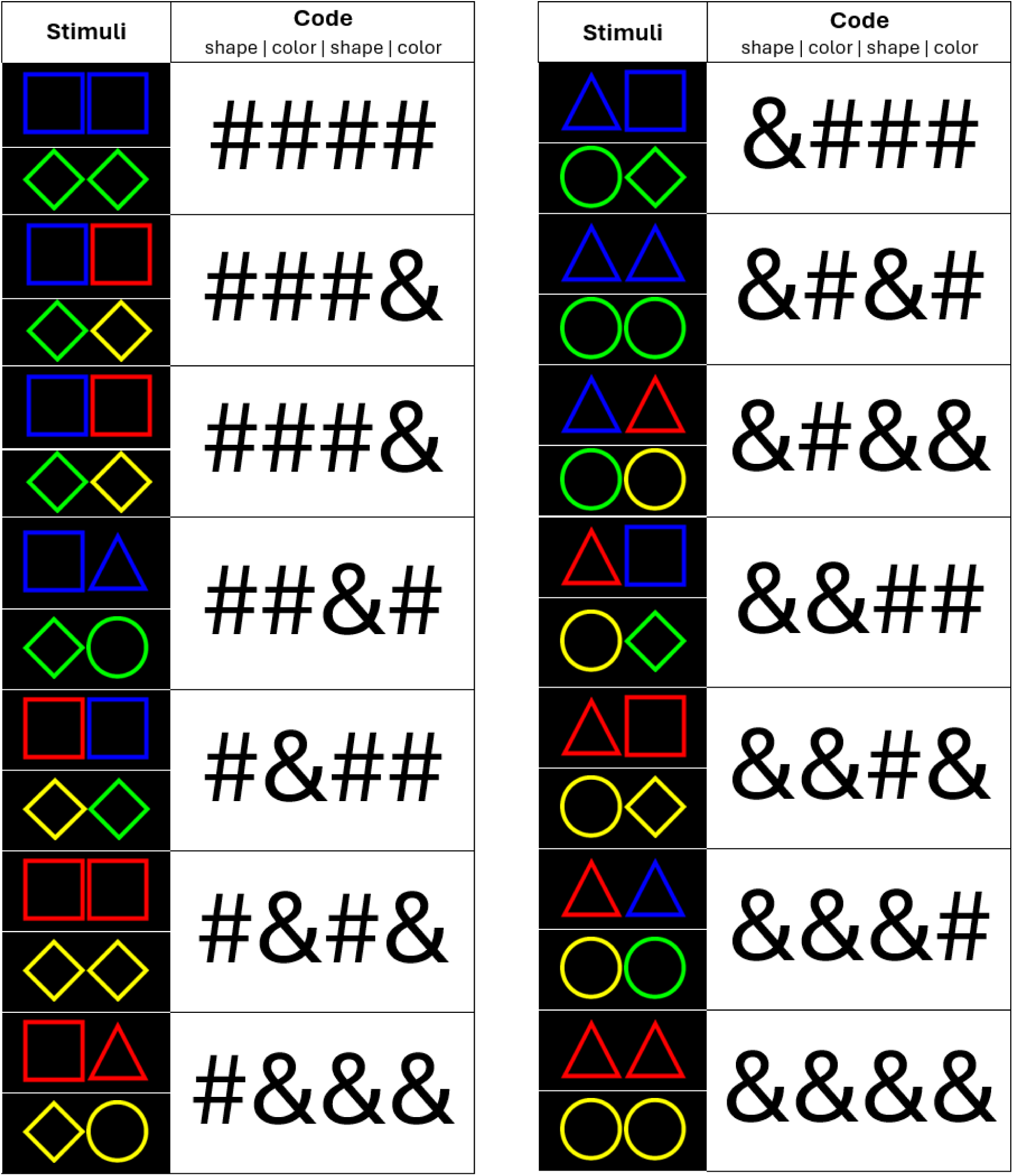
Stimuli for task “catch” trials.

**Table S2.** Reaction time analysis – main fMRI task Results of the Reaction Time (RT) analysis of the behavioral data during the fMRI task. Factors were sequence step (1, 2, 3, 4), planning duration (short, medium, long), and their interaction.

|  | Sum Sq | Mean Sq | NumDF | DenDF | F value | Pr(>F) |
| --- | --- | --- | --- | --- | --- | --- |
| step | 1418.81 | 472.94 | 3 | 32907 | 24256.561 | < 2.2e-16 |
| plan duration | 1.47 | 0.73 | 2 | 32908 | 37.644 | < 2.2e-16 |
| step:plan duration | 6.51 | 1.09 | 6 | 32907 | 55.691 | < 2.2e-16 |

**Table S3.** Post-hoc pairwise differences – main fMRI task Subsequent post-hoc (Tukey HSD) tests performed to identify the direction of significant effects and interactions from the RT analysis of the behavioral data during the fMRI task.

| contrast | estimate | SE | df | t.ratio | p.value |
| --- | --- | --- | --- | --- | --- |
| **Main effect: sequence step** |  |  |  |  |  |
| step1 - step2 | 0.45770 | 0.00218 | 32907 | 210.402 | <.0001 |
| step1 - step3 | 0.46169 | 0.00218 | 32907 | 212.236 | <.0001 |
| step1 - step4 | 0.51072 | 0.00218 | 32907 | 234.778 | <.0001 |
| step2 - step3 | 0.00399 | 0.00218 | 32907 | 1.833 | 0.2576 |
| step2 - step4 | 0.05302 | 0.00218 | 32907 | 24.375 | <.0001 |
| step3 - step4 | 0.04904 | 0.00218 | 32907 | 22.542 | <.0001 |
| **Main effect: plan duration** |  |  |  |  |  |
| long - med | -0.00685 | 0.00189 | 32907 | -3.625 | 0.0008 |
| long - short | -0.01629 | 0.00189 | 32908 | -8.633 | <.0001 |
| med - short | -0.00944 | 0.00189 | 32908 | -5.034 | <.0001 |
| **Interaction: sequence step X plan duration** |  |  |  |  |  |
| step = 1: |  |  |  |  |  |
| long - med | -0.031884 | 0.00378 | 32907 | -8.434 | <.0001 |
| long - short | -0.075597 | 0.00377 | 32907 | -20.032 | <.0001 |
| med - short | -0.043713 | 0.00375 | 32907 | -11.658 | <.0001 |
| step = 2: |  |  |  |  |  |
| long - med | 0.003342 | 0.00378 | 32907 | 0.884 | 0.6504 |
| long - short | 0.007083 | 0.00377 | 32907 | 1.877 | 0.1454 |
| med - short | 0.003741 | 0.00375 | 32907 | 0.998 | 0.5783 |
| step = 3: |  |  |  |  |  |
| long - med | 0.001336 | 0.00378 | 32907 | 0.353 | 0.9334 |
| long - short | 0.002837 | 0.00377 | 32907 | 0.752 | 0.7325 |
| med - short | 0.001501 | 0.00375 | 32907 | 0.400 | 0.9155 |
| step = 4: |  |  |  |  |  |
| long - med | -0.000210 | 0.00378 | 32907 | -0.056 | 0.9983 |
| long - short | 0.000508 | 0.00377 | 32907 | 0.135 | 0.9901 |
| med - short | 0.000718 | 0.00375 | 32907 | 0.191 | 0.9800 |

**Table S4.** Activation value maps for the Multivariate Pattern Analyses. Activation values for key brain regions (clusters) identified during multivariate analyses: searchlight cross-decoding from planning to execution, and searchlight RSA comparing all 4 conditions during the planning phase.

| Region | voxels | Peak<br>t | MNI coords<br>(peak) |  |  | Cluster<br>p |
| --- | --- | --- | --- | --- | --- | --- |
|  |  |  | x | y | z |  |
| Searchlight-cross-decoding<br>planning -> execution<br>(abstract) |  |  |  |  |  |  |
| Lenticular nucleus, Pallidum, left<br>Lenticular nucleus, Putamen, left | 29 | 5.49 | -26 | -14 | -4 | 0.071<br>(uncorr.) |
| Superior frontal gyrus, medial, right<br>Medial frontal gyrus, right | 26 | 4.22 | 10 | 46 | 40 | 0.098<br>(uncorr.) |
| Middle frontal gyrus, left<br>Inferior frontal gyrus, triangular part, left | 24 | 3.94<br>3.60 | -48<br>-42 | 30<br>36 | 30<br>24 | 0.098<br>(uncorr.) |
| Searchlight-cross-decoding<br>planning -> execution<br>(perceptual) |  |  |  |  |  |  |
| Calcarine fissure and surrounding cortex, right<br>Calcarine fissure and surrounding cortex, left | 107 | 4.80<br>4.04 | 4<br>-10 | -76<br>-76 | 10<br>10 | 0.046<br>(uncorr.) |
| Searchlight-RSA, planning<br>(abstract) |  |  |  |  |  |  |
| Superior frontal gyrus, medial, right<br>Medial frontal gyrus, right<br>Superior frontal gyrus, medial, left<br>Medial frontal gyrus, left | 46 | 4.27<br>3.91<br>4.23 | 3<br>6<br>-3 | 38<br>47<br>50 | 44<br>44<br>44 | 0.014<br>(FWE) |
| Searchlight-RSA, planning<br>(perceptual) |  |  |  |  |  |  |
| Middle occipital gyrus, left<br>Superior occipital gyrus, left<br>Inferior occipital gyrus, left<br>Calcarine fissure and surrounding cortex, left<br>Lingual gyrus, left<br>Fusiform gyrus, left | 120 | 6.12<br>4.66 | -18<br>-12 | -91<br>-97 | 1<br>11 | 0.000<br>(FWE) |

**Figure S1.**
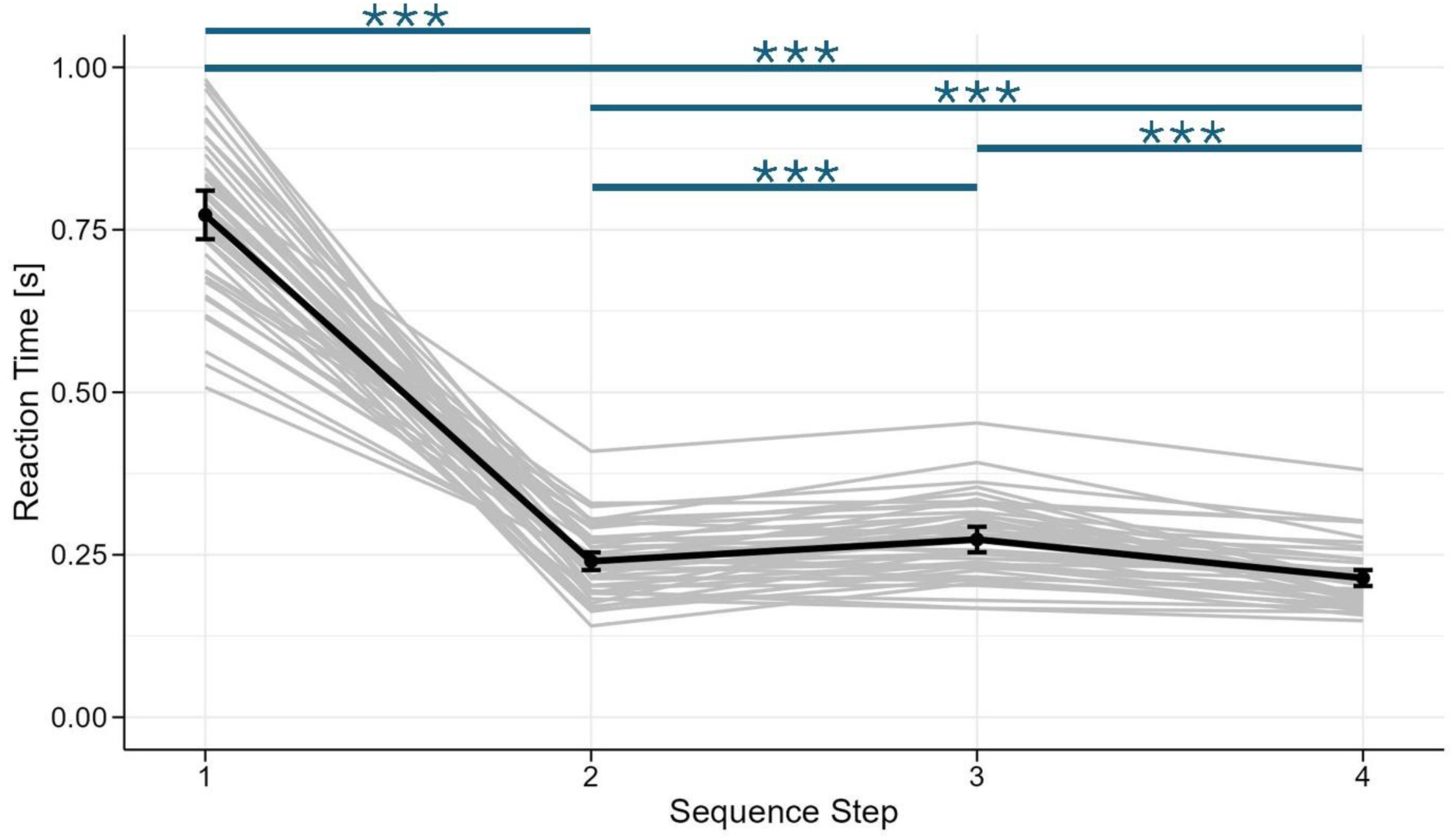
Reaction time (RT) results – behavioral training. Horizontal lines represent post-hoc pairwise comparisons following the main effect of sequence step on RT. Bold black line represents the average RT across participants, gray lines represent individual participants. Asterisks denote statistical significance at p < 0.0001.

**Figure S2.**
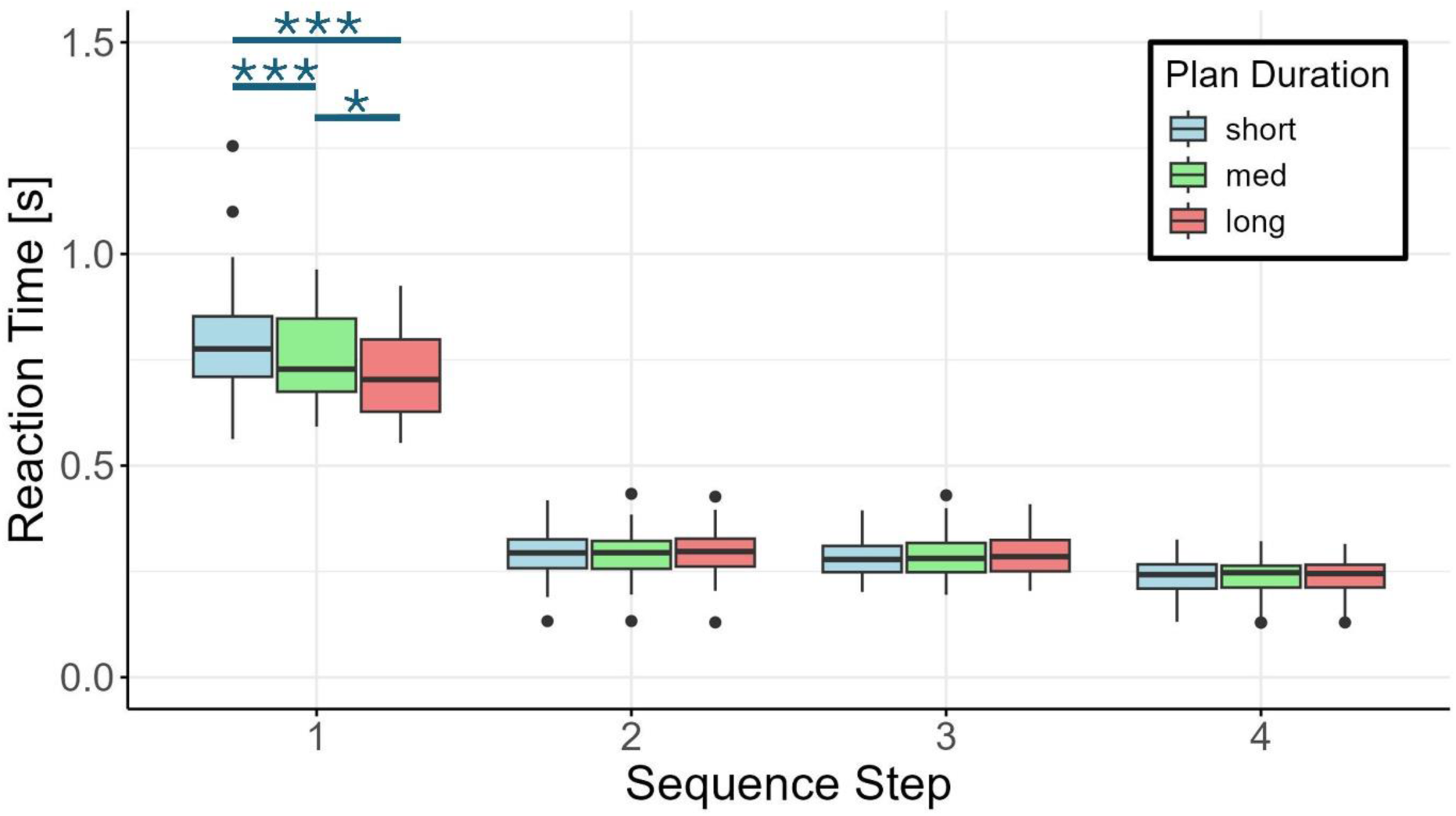
Post-hoc pairwise difference for the main effect of plan duration – main fMRI task.

**Figure S3.**
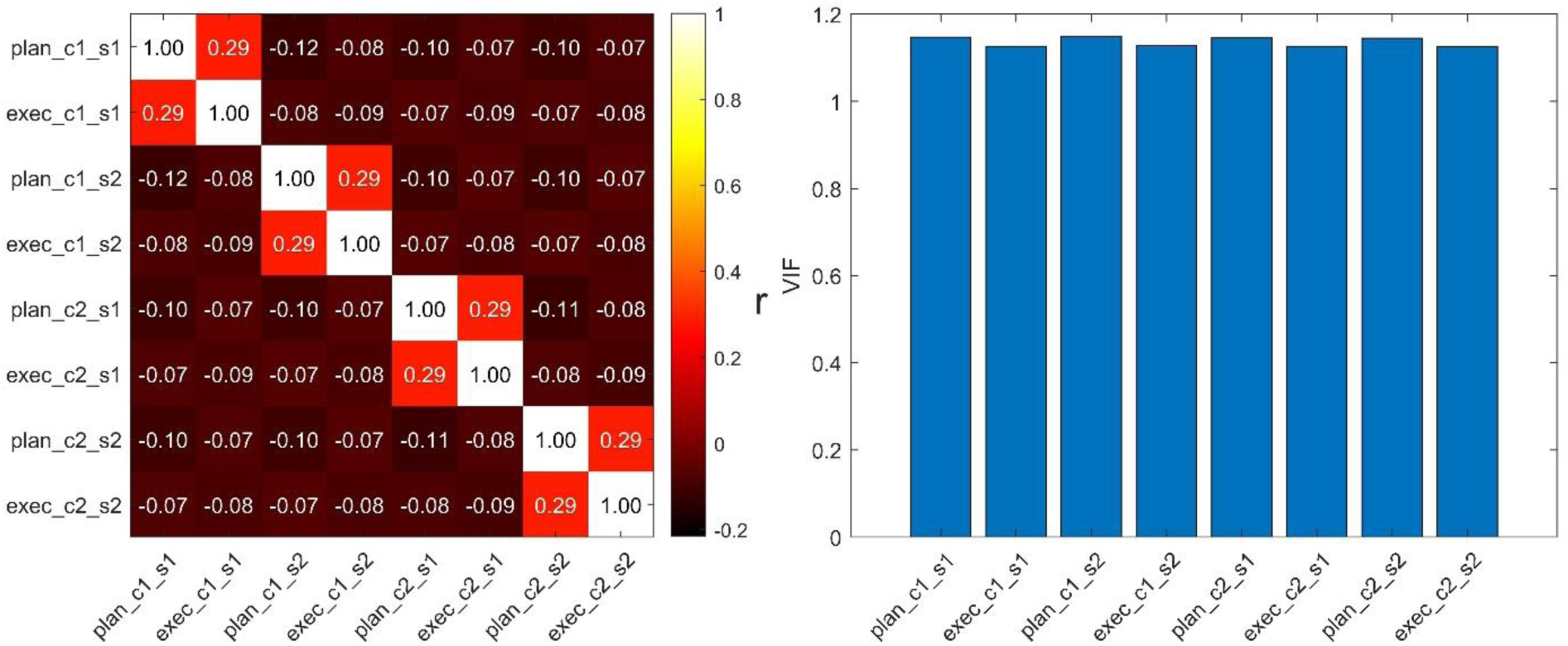
Collinearity assessment: correlation between regressors of interest Our primary analysis relied on training a classifier on activations from the planning phase and testing it on activity from the execution phase. To evaluate that these regressors were not problematically correlated and sufficiently independent for a successful cross-decoding analysis, we evaluated the collinearity in our regressors of interest. For each subject, we extracted the filtered and whitened design matrix from their GLM specification (see main text methods). Within each experimental run, we computed two metrics for the eight regressors of interest (four planning and four execution regressors for conditions c1s1, c1s2, c2s1, c2s2, respectively): a full regressor (8×8) correlation matrix (left panel), from which we could assess the relationship of each regressor to the rest, and the variance inflation factor (VIF (Mumford et al., 2025)) for each regressor (right panel). The VIF of regressor *X*_*i*_ is defined as *VIF*_*i*_ = 1⁄(1 − *R*_i_^2^), where *R*_i_^2^ is the coefficient of determination obtained by regressing *X*_*i*_ on all other regressors in the model (*X*_*i*_ = β_1_*X*_1_ + ⋯ + β_*i*−1_*X*_*i*−1_ + β_*i*+1_*X*_*i*+1_ + ⋯ + β_*N*_*X*_*N*_ + ε) (Mumford et al., 2015, 2025). The VIF provides a direct measure of collinearity since its value indicates the degree at which one regressor represents a linear combination of the other regressors (Mumford et al., 2015), a property that pairwise correlations might not always capture. Higher values of VIF denote higher presence of collinearity, with values greater than 5 or 10 considered problematic (Mumford et al., 2015) (but see (O’brien, 2007)). These two metrics were calculated at the run level before being averaged within each subject to create stable per-subject and group-level estimates. As expected, the VIF was low for all regressors of interest (mean = 1.14, SD = 0.03), the correlations were moderate between planning and execution regressors from the same condition (mean = 0.29, SD = 0.04) and low between planning and execution regressors from different conditions (mean = −0.07, SD = 0.017).

**Figure S4.**
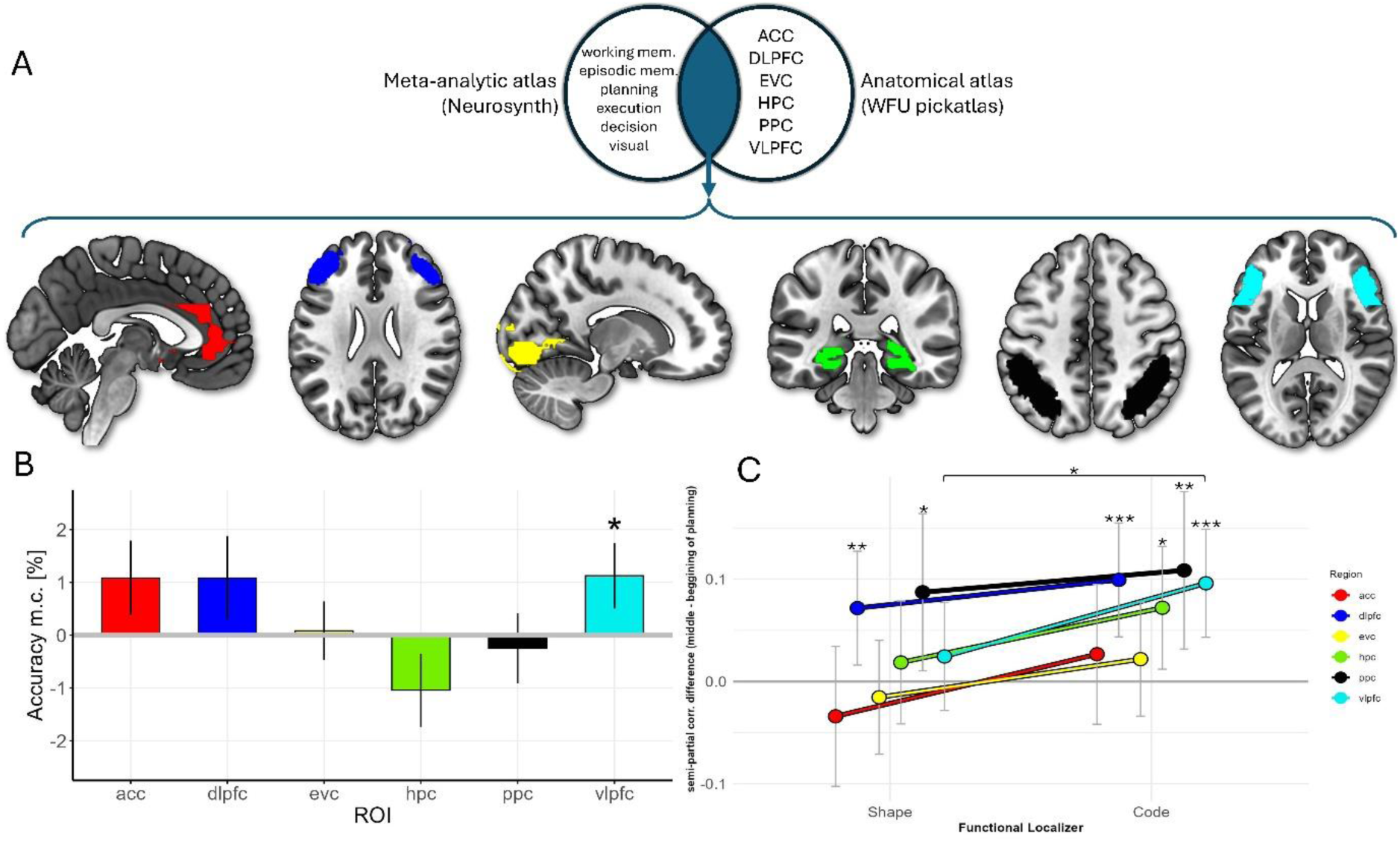
ROI analyses. **A. ROI selection.** As pre-registered, we ran follow-up analyses in specific regions of interest (ROI) for two reasons: 1) our main hypothesis specified that a higher-order context will be represented in planning-related regions, and 2) previous literature has shown that decoding analyses tend to be relatively stringent, especially with regards to information in prefrontal cortex (Bhandari et al., 2018). We selected planning-related ROIs located in: (rostral) anterior cingulate cortex (ACC), ventrolateral prefrontal cortex (VLPFC), dorsolateral prefrontal cortex (DLPFC), posterior parietal cortex (PPC), hippocampus (HPC), and early visual cortex (EVC) as control region. First, a binary mask of each anatomical region was defined, extracted, and combined using the WFU pickatlas software (Maldjian et al., 2003) (version 3.0.5). Second, from the metanalytical tool Neurosynth (https://neurosynth.org/analyses/), we extracted the resulting (FDR-corrected) masks from the search terms “working memory”, “episodic memory”, “planning”, “execution”, “decision”, and “visual.” These metaanalytically-defined masks were binarized and combined into one single aggregate mask. Finally, each anatomically-defined ROI mask was intersected with the aggregate mask, resulting in 6 ROIs. Each ROI mask was registered back to each subject’s native space via estimation of inverse deformation fields. **B. ROI-decoding results.** Decoding was significantly above chance for VLPFC. **C. Semi-partial correlation results.** Abstract representations increased significantly from the beginning to the middle of planning in DLPFC, HPC, PPC, and VLPFC. Perceptual representations increased significantly from the beginning to the middle of planning in DLPFC and PPC. The dynamics between abstract and perceptual representations were significantly different only for VLPFC. ROI-RSA results were not significant in any of the ROIs.

## REFERENCES

Abe, M., Hanakawa, T., Takayama, Y., Kuroki, C., Ogawa, S., & Fukuyama, H. (2007). Functional coupling of human prefrontal and premotor areas during cognitive manipulation. The Journal of Neuroscience: The Official Journal of the Society for Neuroscience, 27(13), 3429–3438. 10.1523/JNEUROSCI.4273-06.2007

Afshar, A., Santhanam, G., Yu, B. M., Ryu, S. I., Sahani, M., & Shenoy, K. V. (2011). Single-Trial Neural Correlates of Arm Movement Preparation. Neuron, 71(3), 555–564. 10.1016/j.neuron.2011.05.047

Alejandro, R. J., & Holroyd, C. B. (2024). Hierarchical control over foraging behavior by anterior cingulate cortex. Neuroscience & Biobehavioral Reviews, 160, 105623. 10.1016/j.neubiorev.2024.105623

Alexander, W. H., & Brown, J. W. (2015). Hierarchical Error Representation: A Computational Model of Anterior Cingulate and Dorsolateral Prefrontal Cortex. Neural Computation, 27(11), 2354–2410. 10.1162/NECO_a_00779

Ariani, G., Oosterhof, N. N., & Lingnau, A. (2018). Time-resolved decoding of planned delayed and immediate prehension movements. Cortex, 99, 330–345. 10.1016/j.cortex.2017.12.007

Avants, B., Epstein, C., Grossman, M., & Gee, J. (2008). Symmetric diffeomorphic image registration with cross-correlation: Evaluating automated labeling of elderly and neurodegenerative brain. Medical Image Analysis, 12(1), 26–41. 10.1016/j.media.2007.06.004

Badre, D. (2025). Cognitive Control. Annual Review of Psychology, 76(1), 167–195. 10.1146/annurev-psych-022024-103901

Balaguer, J., Spiers, H., Hassabis, D., & Summerfield, C. (2016). Neural Mechanisms of Hierarchical Planning in a Virtual Subway Network. Neuron, 90(4), 893–903. 10.1016/j.neuron.2016.03.037

Behzadi, Y., Restom, K., Liau, J., & Liu, T. T. (2007). A component based noise correction method (CompCor) for BOLD and perfusion based fMRI. NeuroImage, 37(1), 90–101. 10.1016/j.neuroimage.2007.04.042

Bernardi, S., Benna, M. K., Rigotti, M., Munuera, J., Fusi, S., & Salzman, C. D. (2020). The Geometry of Abstraction in the Hippocampus and Prefrontal Cortex. Cell, 183(4), 954–967.e21. 10.1016/j.cell.2020.09.031

Botvinick, M. M. (2008). Hierarchical models of behavior and prefrontal function. Trends in Cognitive Sciences, 12(5), 201–208. 10.1016/j.tics.2008.02.009

Brandi, M.-L., Wohlschläger, A., Sorg, C., & Hermsdörfer, J. (2014). The Neural Correlates of Planning and Executing Actual Tool Use. The Journal of Neuroscience, 34(39), 13183–13194. 10.1523/JNEUROSCI.0597-14.2014

Brunec, I. K., & Momennejad, I. (2022). Predictive Representations in Hippocampal and Prefrontal Hierarchies. The Journal of Neuroscience, 42(2), 299–312. 10.1523/JNEUROSCI.1327-21.2021

Bunge, S. A., Hazeltine, E., Scanlon, M. D., Rosen, A. C., & Gabrieli, J. D. E. (2002). Dissociable Contributions of Prefrontal and Parietal Cortices to Response Selection. NeuroImage, 17(3), 1562–1571. 10.1006/nimg.2002.1252

Chatham, C. H., & Badre, D. (2015). Multiple gates on working memory. Current Opinion in Behavioral Sciences, 1, 23–31. 10.1016/j.cobeha.2014.08.001

Chikazoe, J., Jimura, K., Hirose, S., Yamashita, K., Miyashita, Y., & Konishi, S. (2009). Preparation to Inhibit a Response Complements Response Inhibition during Performance of a Stop-Signal Task. The Journal of Neuroscience, 29(50), 15870–15877. 10.1523/JNEUROSCI.3645-09.2009

Cisek, P., & Kalaska, J. F. (2010). Neural Mechanisms for Interacting with a World Full of Action Choices. Annual Review of Neuroscience, 33(1), 269–298. 10.1146/annurev.neuro.051508.135409

Cohen, J. D., Perlstein, W. M., Braver, T. S., Nystrom, L. E., Noll, D. C., Jonides, J., & Smith, E. E. (1997). Temporal dynamics of brain activation during a working memory task. Nature, 386(6625), 604–608. 10.1038/386604a0

Cohen, M. X., & Frank, M. J. (2009). Neurocomputational models of basal ganglia function in learning, memory and choice. Behavioural Brain Research, 199(1), 141–156. 10.1016/j.bbr.2008.09.029

Cole, M. W., Bagic, A., Kass, R., & Schneider, W. (2010). Prefrontal Dynamics Underlying Rapid Instructed Task Learning Reverse with Practice. The Journal of Neuroscience, 30(42), 14245–14254. 10.1523/jneurosci.1662-10.2010

Coles, M. G. H., Gratton, G., Bashore, T. R., Eriksen, C. W., & Donchin, E. (1985). A psychophysiological investigation of the continuous flow model of human information processing. Journal of Experimental Psychology: Human Perception and Performance, 11(5), 529–553. 10.1037/0096-1523.11.5.529

Colin, T. R., Ikink, I., & Holroyd, C. B. (2025). Distributed Representations for Cognitive Control in Frontal Medial Cortex. Journal of Cognitive Neuroscience, 37(5), 941–969. 10.1162/jocn_a_02285

Collins, A. G. E., & Frank, M. J. (2013). Cognitive control over learning: Creating, clustering, and generalizing task-set structure. Psychological Review, 120(1), 190–229. 10.1037/a0030852

Crescentini, C., Seyed-Allaei, S., Vallesi, A., & Shallice, T. (2012). Two networks involved in producing and realizing plans. Neuropsychologia, 50(7), 1521–1535. 10.1016/j.neuropsychologia.2012.03.005

Curtis, C. E., & D’Esposito, M. (2003). Persistent activity in the prefrontal cortex during working memory. Trends in Cognitive Sciences, 7(9), 415–423. 10.1016/S1364-6613(03)00197-9

Dale, A. M. (1999). Optimal experimental design for event-related fMRI. Human Brain Mapping, 8(2–3), 109–114. 10.1002/(SICI)1097-0193(1999)8:2/3%253C109::AID-HBM7%253E3.0.CO;2-W

Dehaene, S., & Changeux, J.-P. (1997). A hierarchical neuronal network for planning behavior. Proceedings of the National Academy of Sciences, 94(24), 13293–13298. 10.1073/pnas.94.24.13293

Dolfen, N., Reverberi, S., Op De Beeck, H., King, B. R., & Albouy, G. (2024). The Hippocampus Represents Information about Movements in Their Temporal Position in a Learned Motor Sequence. The Journal of Neuroscience, 44(37), e0584242024. 10.1523/JNEUROSCI.0584-24.2024

Duverne, S., & Koechlin, E. (2017). Hierarchical Control of Behaviour in Human Prefrontal Cortex. In T. Egner (Ed.), The Wiley Handbook of Cognitive Control (1st ed., pp. 207–220). Wiley. 10.1002/9781118920497.ch12

Eckstein, M. K., & Collins, A. G. (2021). How the Mind Creates Structure: Hierarchical Learning of Action Sequences. Proceedings of the Annual Meeting of the Cognitive Science Society. Annual Meeting of the Cognitive Science Society. https://escholarship.org/uc/item/4v65s77x

Elsinger, C. L., Harrington, D. L., & Rao, S. M. (2006). From preparation to online control: Reappraisal of neural circuitry mediating internally generated and externally guided actions. NeuroImage, 31(3), 1177–1187. 10.1016/j.neuroimage.2006.01.041

Fermin, A. S. R., Yoshida, T., Yoshimoto, J., Ito, M., Tanaka, S. C., & Doya, K. (2016). Model-based action planning involves cortico-cerebellar and basal ganglia networks. Scientific Reports, 6(1), 31378. 10.1038/srep31378

Fincham, J. M., Carter, C. S., Van Veen, V., Stenger, V. A., & Anderson, J. R. (2002). Neural mechanisms of planning: A computational analysis using event-related fMRI. Proceedings of the National Academy of Sciences, 99(5), 3346–3351. 10.1073/pnas.052703399

Foinikianaki, E., Ikink, I., Colin, T. R., Alejandro, R. J., & Holroyd, C. B. (2025). ACC representations of reward-driven motivation over hierarchically-organized behavior. NeuroImage, 317, 121380. 10.1016/j.neuroimage.2025.121380

Fuster, J. M., & Alexander, G. E. (1971). Neuron Activity Related to Short-Term Memory. Science, 173(3997), 652–654. 10.1126/science.173.3997.652

Gallivan, J. P., McLean, D. A., Valyear, K. F., Pettypiece, C. E., & Culham, J. C. (2011). Decoding Action Intentions from Preparatory Brain Activity in Human Parieto-Frontal Networks. Journal of Neuroscience, 31(26), 9599–9610. 10.1523/JNEUROSCI.0080-11.2011

Gorgolewski, K., Burns, C. D., Madison, C., Clark, D., Halchenko, Y. O., Waskom, M. L., & Ghosh, S. S. (2011). Nipype: A Flexible, Lightweight and Extensible Neuroimaging Data Processing Framework in Python. Frontiers in Neuroinformatics, 5. 10.3389/fninf.2011.00013

Gorgolewski, K. J., Alfaro-Almagro, F., Auer, T., Bellec, P., Capotă, M., Chakravarty, M. M., Churchill, N. W., Cohen, A. L., Craddock, R. C., Devenyi, G. A., Eklund, A., Esteban, O., Flandin, G., Ghosh, S. S., Guntupalli, J. S., Jenkinson, M., Keshavan, A., Kiar, G., Liem, F., … Poldrack, R. A. (2017). BIDS apps: Improving ease of use, accessibility, and reproducibility of neuroimaging data analysis methods. PLOS Computational Biology, 13(3), e1005209. 10.1371/journal.pcbi.1005209

Graybiel, A. M. (2000). The basal ganglia. Current Biology, 10(14), R509–R511. 10.1016/S0960-9822(00)00593-5

Greve, D. N., & Fischl, B. (2009). Accurate and robust brain image alignment using boundary-based registration. NeuroImage, 48(1), 63–72. 10.1016/j.neuroimage.2009.06.060

Groenewegen, H. J. (2003). The Basal Ganglia and Motor Control. Neural Plasticity, 10(1–2), 107–120. 10.1155/NP.2003.107

Grossman, C. D., Man, V., & O’Doherty, J. P. (2025). The representation and valuation of subgoals in the human brain during model-based hierarchical behavior. 10.1101/2025.03.24.645084

Hall-McMaster, S., Tomov, M. S., Gershman, S. J., & Schuck, N. W. (2025). Neural evidence that humans reuse strategies to solve new tasks. PLOS Biology, 23(6), e3003174. 10.1371/journal.pbio.3003174

Hanakawa, T., Dimyan, M. A., & Hallett, M. (2008). Motor Planning, Imagery, and Execution in the Distributed Motor Network: A Time-Course Study with Functional MRI. Cerebral Cortex, 18(12), 2775–2788. 10.1093/cercor/bhn036

Hebart, M. N., Görgen, K., & Haynes, J.-D. (2015). The Decoding Toolbox (TDT): A versatile software package for multivariate analyses of functional imaging data. Frontiers in Neuroinformatics, 8. 10.3389/fninf.2014.00088

Henderson, M. M., Rademaker, R. L., & Serences, J. T. (2022). Flexible utilization of spatial- and motor-based codes for the storage of visuo-spatial information. eLife, 11, e75688. 10.7554/eLife.75688

Ho, M. K., Abel, D., Correa, C. G., Littman, M. L., Cohen, J. D., & Griffiths, T. L. (2022). People construct simplified mental representations to plan. Nature, 606(7912), 129–136. 10.1038/s41586-022-04743-9

Holroyd, C. B., Ribas-Fernandes, J. J. F., Shahnazian, D., Silvetti, M., & Verguts, T. (2018). Human midcingulate cortex encodes distributed representations of task progress. Proceedings of the National Academy of Sciences, 115(25), 6398–6403. 10.1073/pnas.1803650115

Holroyd, C. B., & Verguts, T. (2021). The Best Laid Plans: Computational Principles of Anterior Cingulate Cortex. Trends in Cognitive Sciences, 25(4), 316–329. 10.1016/j.tics.2021.01.008

Holroyd, C. B., & Yeung, N. (2012). Motivation of extended behaviors by anterior cingulate cortex. Trends in Cognitive Sciences, 16(2), 122–128. 10.1016/j.tics.2011.12.008

Hommel, B., Müsseler, J., Aschersleben, G., & Prinz, W. (2001). The Theory of Event Coding (TEC): A framework for perception and action planning. Behavioral and Brain Sciences, 24(5), 849–878. 10.1017/S0140525X01000103

Hoshi, E., & Tanji, J. (2004). Area-Selective Neuronal Activity in the Dorsolateral Prefrontal Cortex for Information Retrieval and Action Planning. Journal of Neurophysiology, 91(6), 2707–2722. 10.1152/jn.00904.2003

Huettel, S. A. (2012). Event-related fMRI in cognition. NeuroImage, 62(2), 1152–1156. 10.1016/j.neuroimage.2011.08.113

Ito, M. (2008). Control of mental activities by internal models in the cerebellum. Nature Reviews Neuroscience, 9(4), 304–313. 10.1038/nrn2332

James, G., Witten, D., Hastie, T., & Tibshirani, R. (2013). An Introduction to Statistical Learning (Vol. 103). Springer New York. 10.1007/978-1-4614-7138-7

Jenkinson, M., Bannister, P., Brady, M., & Smith, S. (2002). Improved Optimization for the Robust and Accurate Linear Registration and Motion Correction of Brain Images. NeuroImage, 17(2), 825–841. 10.1006/nimg.2002.1132

Josephs, O., Turner, R., & Friston, K. (1997). Event-related f MRI. Human Brain Mapping, 5(4), 243–248. 10.1002/(SICI)1097-0193(1997)5:4%253C243::AID-HBM7%253E3.0.CO;2-3

Kawagoe, R., Takikawa, Y., & Hikosaka, O. (1998). Expectation of reward modulates cognitive signals in the basal ganglia. Nature Neuroscience, 1(5), 411–416. 10.1038/1625

Koechlin, E., Corrado, G., Pietrini, P., & Grafman, J. (2000). Dissociating the role of the medial and lateral anterior prefrontal cortex in human planning. Proceedings of the National Academy of Sciences, 97(13), 7651–7656. 10.1073/pnas.130177397

Konidaris, G., Kaelbling, L. P., & Lozano-Perez, T. (2018). From Skills to Symbols: Learning Symbolic Representations for Abstract High-Level Planning. Journal of Artificial Intelligence Research, 61, 215–289. 10.1613/jair.5575

Kriegeskorte, N., Goebel, R., & Bandettini, P. (2006). Information-based functional brain mapping. Proceedings of the National Academy of Sciences, 103(10), 3863–3868. 10.1073/pnas.0600244103

Kriegeskorte, N., Mur, M., & Bandettini, P. (2008). Representational similarity analysis – connecting the branches of systems neuroscience. Frontiers in Systems Neuroscience. 10.3389/neuro.06.004.2008

Kruggel, F., & Von Cramon, D. Y. (1999). Temporal properties of the hemodynamic response in functional MRI. Human Brain Mapping, 8(4), 259–271. 10.1002/(SICI)1097-0193(1999)8:4%253C259::AID-HBM9%253E3.0.CO;2-K

Kruskal, J. B. (1964). Nonmetric Multidimensional Scaling: A Numerical Method. Psychometrika, 29(2), 115–129. 10.1007/BF02289694

Kubota, K., & Niki, H. (1971). Prefrontal cortical unit activity and delayed alternation performance in monkeys. Journal of Neurophysiology, 34(3), 337–347. 10.1152/jn.1971.34.3.337

Lazeron, R. H. C., Rombouts, S. A. R. B., Machielsen, W. C. M., Scheltens, P., Witter, M. P., Uylings, H. B. M., & Barkhof, F. (2000). Visualizing Brain Activation during Planning: The Tower of London Test Adapted for Functional MR Imaging. American Journal of Neuroradiology, 21(8), 1407.

Lee, W.-T., Hazeltine, E., & Jiang, J. (2024). Decoding task representations that support generalization in hierarchical task. 10.1101/2024.12.02.626403

Liang, Q., Li, J., Zheng, S., Liao, J., & Huang, R. (2022). Dynamic Causal Modelling of Hierarchical Planning. NeuroImage, 258, 119384. 10.1016/j.neuroimage.2022.119384

Mattar, M. G., & Lengyel, M. (2022). Planning in the brain. Neuron, 110(6), 914–934. 10.1016/j.neuron.2021.12.018

Mian, M. K., Sheth, S. A., Patel, S. R., Spiliopoulos, K., Eskandar, E. N., & Williams, Z. M. (2014). Encoding of rules by neurons in the human dorsolateral prefrontal cortex. Cerebral Cortex, 24(3), 807–816. 10.1093/cercor/bhs361

Miller, E. K., & Buschman, T. J. (2008). Rules through recursion: How interactions between the frontal cortex and basal ganglia may build abstract, complex rules from concrete, simple ones. Neuroscience of Rule-Guided Behavior, 419–440.

Miller, G. A., Galanter, E., & Pribram, K. H. (1960). Plans and the structure of behavior. Henry Holt and Co. 10.1037/10039-000

Momennejad, I., & Haynes, J.-D. (2012). Human anterior prefrontal cortex encodes the ‘what’ and ‘when’ of future intentions. NeuroImage, 61(1), 139–148. 10.1016/j.neuroimage.2012.02.079

Momennejad, I., & Haynes, J.-D. (2013). Encoding of Prospective Tasks in the Human Prefrontal Cortex under Varying Task Loads. The Journal of Neuroscience, 33(44), 17342–17349. 10.1523/JNEUROSCI.0492-13.2013

Mumford, J. A., Demidenko, M. I., Bjork, J. M., Chaarani, B., Feczko, E. J., Garavan, H. P., Hagler Jr., D. J., Nelson, S. M., Wager, T. D., & Poldrack, R. A. (2025). Unintended bias in the pursuit of collinearity solutions in fMRI analysis. Imaging Neuroscience, 3, IMAG.a.958. 10.1162/IMAG.a.958

Mumford, J. A., Poline, J.-B., & Poldrack, R. A. (2015). Orthogonalization of Regressors in fMRI Models. PLOS ONE, 10(4), e0126255. 10.1371/journal.pone.0126255

Nili, H., Wingfield, C., Walther, A., Su, L., Marslen-Wilson, W., & Kriegeskorte, N. (2014). A Toolbox for Representational Similarity Analysis. PLoS Computational Biology, 10(4), e1003553. 10.1371/journal.pcbi.1003553

Norman, K. A., Polyn, S. M., Detre, G. J., & Haxby, J. V. (2006). Beyond mind-reading: Multi-voxel pattern analysis of fMRI data. Trends in Cognitive Sciences, 10(9), 424–430. 10.1016/j.tics.2006.07.005

O’brien, R. M. (2007). A Caution Regarding Rules of Thumb for Variance Inflation Factors. Quality & Quantity, 41(5), 673–690. 10.1007/s11135-006-9018-6

Palenciano, A. F., Senoussi, M., Formica, S., & González-García, C. (2023). Canonical template tracking: Measuring the activation state of specific neural representations. Frontiers in Neuroimaging, 1, 974927. 10.3389/fnimg.2022.974927

Papitto, G., Friederici, A. D., & Zaccarella, E. (2024). Distinct neural mechanisms for action access and execution in the human brain: Insights from an fMRI study. Cerebral Cortex, 34(4), bhae163. 10.1093/cercor/bhae163

Patriat, R., Reynolds, R. C., & Birn, R. M. (2017). An improved model of motion-related signal changes in fMRI. NeuroImage, 144, 74–82. 10.1016/j.neuroimage.2016.08.051

Pochon, J.-B. (2001). The Role of Dorsolateral Prefrontal Cortex in the Preparation of Forthcoming Actions: An fMRI Study. Cerebral Cortex, 11(3), 260–266. 10.1093/cercor/11.3.260

Power, J. D., Mitra, A., Laumann, T. O., Snyder, A. Z., Schlaggar, B. L., & Petersen, S. E. (2014). Methods to detect, characterize, and remove motion artifact in resting state fMRI. NeuroImage, 84, 320–341. 10.1016/j.neuroimage.2013.08.048

Reverberi, C., Görgen, K., & Haynes, J.-D. (2012). Compositionality of Rule Representations in Human Prefrontal Cortex. Cerebral Cortex, 22(6), 1237–1246. 10.1093/cercor/bhr200

Rosenbaum, D. A. (1980). Human movement initiation: Specification of arm, direction, and extent. Journal of Experimental Psychology: General, 109(4), 444–474. 10.1037/0096-3445.109.4.444

Rosenbaum, D. A., Kenny, S. B., & Derr, M. A. (1983). Hierarchical control of rapid movement sequences. Journal of Experimental Psychology: Human Perception and Performance, 9(1), 86–102. 10.1037/0096-1523.9.1.86

Rousseau, C., Barbiero, M., Pozzo, T., Papaxanthis, C., & White, O. (2021). Actual and Imagined Movements Reveal a Dual Role of the Insular Cortex for Motor Control. Cerebral Cortex, 31(5), 2586–2594. 10.1093/cercor/bhaa376

Rowe, J. B., Owen, A. M., Johnsrude, I. S., & Passingham, R. E. (2001). Imaging the mental components of a planning task. Neuropsychologia, 39(3), 315–327. 10.1016/S0028-3932(00)00109-3

Ruge, H., Müller, S. C., & Braver, T. S. (2010). Anticipating the consequences of action: An fMRI study of intention-based task preparation: Anticipating the consequences of action. Psychophysiology, no-no. 10.1111/j.1469-8986.2010.01027.x

Sakai, K., & Passingham, R. E. (2003). Prefrontal interactions reflect future task operations. Nature Neuroscience, 6(1), 75–81. 10.1038/nn987

Sakai, K., & Passingham, R. E. (2006). Prefrontal Set Activity Predicts Rule-Specific Neural Processing during Subsequent Cognitive Performance. The Journal of Neuroscience, 26(4), 1211–1218. 10.1523/jneurosci.3887-05.2006

Sakai, K., Rowe, J. B., & Passingham, R. E. (2002). Active maintenance in prefrontal area 46 creates distractor-resistant memory. Nature Neuroscience, 5(5), 479–484. 10.1038/nn846

Samejima, K., Ueda, Y., Doya, K., & Kimura, M. (2005). Representation of Action-Specific Reward Values in the Striatum. Science, 310(5752), 1337–1340. 10.1126/science.1115270

Satterthwaite, T. D., Elliott, M. A., Gerraty, R. T., Ruparel, K., Loughead, J., Calkins, M. E., Eickhoff, S. B., Hakonarson, H., Gur, R. C., Gur, R. E., & Wolf, D. H. (2013). An improved framework for confound regression and filtering for control of motion artifact in the preprocessing of resting-state functional connectivity data. NeuroImage, 64, 240–256. 10.1016/j.neuroimage.2012.08.052

Schumacher, E. H., & D’Esposito, M. (2002). Neural implementation of response selection in humans as revealed by localized effects of stimulus–response compatibility on brain activation. Human Brain Mapping, 17(3), 193–201. 10.1002/hbm.10063

Simon, D. A., & Daw, N. D. (2011). Neural Correlates of Forward Planning in a Spatial Decision Task in Humans. The Journal of Neuroscience, 31(14), 5526–5539. 10.1523/JNEUROSCI.4647-10.2011

Skorstad, J., Genter, D., & Medin, D. (1988). Abstraction Processes during Concept Learning: A Structural view. Proceedings of the Annual Meeting of the Cognitive Science Society, 10. https://escholarship.org/uc/item/9mg8p8hp#author

Tanaka, M., Kunimatsu, J., Suzuki, T. W., Kameda, M., Ohmae, S., Uematsu, A., & Takeya, R. (2021). Roles of the Cerebellum in Motor Preparation and Prediction of Timing. Neuroscience, 462, 220–234. 10.1016/j.neuroscience.2020.04.039

Theves, S., Neville, D. A., Fernández, G., & Doeller, C. F. (2021). Learning and Representation of Hierarchical Concepts in Hippocampus and Prefrontal Cortex. The Journal of Neuroscience, 41(36), 7675–7686. 10.1523/JNEUROSCI.0657-21.2021

Tolman, E. C. (1948). Cognitive maps in rats and men. Psychological Review, 55(4), 189–208. 10.1037/h0061626

Tomov, M. S., Yagati, S., Kumar, A., Yang, W., & Gershman, S. J. (2020). Discovery of hierarchical representations for efficient planning. PLOS Computational Biology, 16(4), e1007594. 10.1371/journal.pcbi.1007594

Tosoni, A., Corbetta, M., Calluso, C., Committeri, G., Pezzulo, G., Romani, G. L., & Galati, G. (2014). Decision and action planning signals in human posterior parietal cortex during delayed perceptual choices. European Journal of Neuroscience, 39(8), 1370–1383. 10.1111/ejn.12511

Tricomi, E., Balleine, B. W., & O’Doherty, J. P. (2009). A specific role for posterior dorsolateral striatum in human habit learning. European Journal of Neuroscience, 29(11), 2225–2232. 10.1111/j.1460-9568.2009.06796.x

Turella, L., Rumiati, R., & Lingnau, A. (2020). Hierarchical Action Encoding Within the Human Brain. Cerebral Cortex, 30(5), 2924–2938. 10.1093/cercor/bhz284

Tustison, N. J., Avants, B. B., Cook, P. A., Yuanjie Zheng Egan, A., Yushkevich, P. A., & Gee, J. C. (2010). N4ITK: Improved N3 Bias Correction. IEEE Transactions on Medical Imaging, 29(6), 1310–1320. 10.1109/TMI.2010.2046908

Unterrainer, J. M., & Owen, A. M. (2006). Planning and problem solving: From neuropsychology to functional neuroimaging. Journal of Physiology-Paris, 99(4–6), 308–317. 10.1016/j.jphysparis.2006.03.014

Vaidya, A. R., & Badre, D. (2022). Abstract task representations for inference and control. *Trends in Cognitive Sciences*, S1364661322000675. 10.1016/j.tics.2022.03.009

Vallesi, A., McIntosh, A. R., Shallice, T., & Stuss, D. T. (2009). When Time Shapes Behavior: fMRI Evidence of Brain Correlates of Temporal Monitoring. Journal of Cognitive Neuroscience, 21(6), 1116–1126. 10.1162/jocn.2009.21098

Van Den Heuvel, O. A., Groenewegen, H. J., Barkhof, F., Lazeron, R. H. C., Van Dyck, R., & Veltman, D. J. (2003). Frontostriatal system in planning complexity: A parametric functional magnetic resonance version of tower of london task. NeuroImage, 18(2), 367–374. 10.1016/S1053-8119(02)00010-1

Wagner, G., Koch, K., Reichenbach, J. R., Sauer, H., & Schlösser, R. G. M. (2006). The special involvement of the rostrolateral prefrontal cortex in planning abilities: An event-related fMRI study with the Tower of London paradigm. Neuropsychologia, 44(12), 2337–2347. 10.1016/j.neuropsychologia.2006.05.014

Wientjes, S., & Holroyd, C. B. (2024). The successor representation subserves hierarchical abstraction for goal-directed behavior. PLOS Computational Biology, 20(2), e1011312. 10.1371/journal.pcbi.1011312

Woo, C.-W., Krishnan, A., & Wager, T. D. (2014). Cluster-extent based thresholding in fMRI analyses: Pitfalls and recommendations. NeuroImage, 91, 412–419. 10.1016/j.neuroimage.2013.12.058

Xia, L., & Collins, A. G. E. (2021). Temporal and state abstractions for efficient learning, transfer, and composition in humans. Psychological Review, 128(4), 643–666. 10.1037/rev0000295

Yewbrey, R., & Kornysheva, K. (2024). The Hippocampus Preorders Movements for Skilled Action Sequences. The Journal of Neuroscience, 44(45), e0832242024. 10.1523/JNEUROSCI.0832-24.2024

Yewbrey, R., Mantziara, M., & Kornysheva, K. (2023). Cortical Patterns Shift from Sequence Feature Separation during Planning to Integration during Motor Execution. The Journal of Neuroscience, 43(10), 1742–1756. 10.1523/JNEUROSCI.1628-22.2023

Ying, L., Collins, K. M., Sharma, P., Colas, C., Zhao, K. I., Weller, A., Tavares, Z., Isola, P., Gershman, S. J., Andreas, J. D., Griffiths, T. L., Chollet, F., Allen, K. R., & Tenenbaum, J. B. (2025). *Assessing Adaptive World Models in Machines with Novel Games* (Version 2). arXiv. 10.48550/ARXIV.2507.12821

Zarr, N., & Brown, J. W. (2016). Hierarchical error representation in medial prefrontal cortex. NeuroImage, 124, 238–247. 10.1016/j.neuroimage.2015.08.063

Zhang, Y., Brady, M., & Smith, S. (2001). Segmentation of brain MR images through a hidden Markov random field model and the expectation-maximization algorithm. IEEE Transactions on Medical Imaging, 20(1), 45–57. 10.1109/42.906424

## SUPPLEMENTARY REFERENCES

Bhandari, A., Gagne, C., & Badre, D. (2018). Just above Chance: Is It Harder to Decode Information from Prefrontal Cortex Hemodynamic Activity Patterns? Journal of Cognitive Neuroscience, 30(10), 1473–1498. 10.1162/jocn_a_01291

Maldjian, J. A., Laurienti, P. J., Kraft, R. A., & Burdette, J. H. (2003). An automated method for neuroanatomic and cytoarchitectonic atlas-based interrogation of fMRI data sets. NeuroImage, 19(3), 1233–1239. 10.1016/S1053-8119(03)00169-1

